# Planar cell polarity in the *C. elegans* embryo emerges by differential retention of aPARs at cell-cell contacts

**DOI:** 10.1101/576777

**Authors:** Priyanka Dutta, Devang Odedra, Christian Pohl

**Affiliations:** Buchmann Institute for Molecular Life Sciences and Institute of Biochemistry II, Medical Faculty, Goethe University, Max-von-Laue-Strasse 15, 60438 Frankfurt (Main), Germany

**Author notes:** contributed equally (Priyanka Dutta, Devang Odedra).

## Abstract

Formation of the anteroposterior and dorsoventral body axis in the *Caenorhabditis elegans* embryo depends on cortical actomyosin flows and advection of polarity determinants. The role of this patterning mechanism in tissue polarization immediately after formation of cell-cell contacts is not fully understood. Here, we demonstrate that planar cell polarity (PCP) is established in the *C. elegans* embryo at the time of left-right (l/r) symmetry breaking. At this stage, centripetal cortical flows asymmetrically and differentially advect anterior polarity determinants (aPARs) PAR-3, PAR-6 and PKC-3 from cell-cell contacts to the medial cortex, which results in their unmixing from apical myosin. Advection generally requires GSK-3 and CDC-42, while advection of PAR-6 specifically depends on the RhoGAP PAC-1. Concurrent asymmetric retention of PAR-3, E-cadherin/HMR-1, PAC-1 and opposing retention of the antagonistic Wnt pathway components APC/APR-1 and Frizzled/MOM-5 at apical cell-cell contacts leads to planar asymmetries. The most obvious mark of PCP, asymmetric retention of PAR-3 at posterior cell-cell contacts on the left side of the embryo, is required for proper cytokinetic cell intercalation. Hence, our data uncover how PCP can be established through Wnt signaling as well as dissociation and planar asymmetric retention of aPARs mediated by distinct Rho GTPases and their regulators.

## Introduction

Gradients in cortical tension can give rise to translocation of the contractile actomyosin network underlying the plasma membrane, a phenomenon called cortical flow (Chalut and Paluch, 2016). During animal development, cortical flow serves as a highly versatile biomechanical actuation system for cellular decision making due to differential spatiotemporal regulation and selective coupling to other cortically localized factors, e.g. polarity determinants, adhesion or signaling complexes. Polarized activation of cortical flow and transient, avidity-driven interactions have been shown to lead to advection of anterior polarity factors (aPARs) PAR-3, PAR-6, and PKC-3, thereby establishing the anteroposterior axis in *C. elegans* (Munro et al., 2004; Goehring et al., 2011; Dickinson et al., 2017; Wang et al., 2017; Mittasch et al., 2018). In addition to patterning the anteroposterior axis, where longitudinal cortical flow is required, dorsoventral and left/right (l/r) patterning in *C. elegans* require rotational cortical flow (Singh and Pohl, 2014; Naganathan et al., 2014; Pohl, 2015; Sugioka and Bowerman, 2018). Rotational flow emerges after formation of cell-cell contacts, where contact-dependent asymmetries determine cortical flow dynamics, which in turn determine spindle orientation through coupling to microtubule dynamics (Sugioka and Bowerman, 2018). These roles of cortical flow in patterning are orthologous in higher organisms, where they have been shown to drive decision making processes in development (Woolner and Papalopulu, 2012; Maitre et al., 2016; Roubinet et al., 2017).

Despite the importance of cortical flows in polarized cell division, we know much less about the role of cortical flow during initiation of apicobasal polarity. In *C. elegans*, apicobasal polarity emerges during the second round of cell divisions. Here, the aPAR polarity determinants PAR-3, PAR-6 and PKC-3 that specified the anterior or somatic part of the embryo become restricted to the apical, contact free surfaces of blastomeres. For one of the apical polarity factors, PAR-6, an active process for its exclusion from basolateral cell-cell contacts has been identified (Anderson et al., 2008; Chan and Nance, 2013). While a RhoGAP, PAC-1, inactivates CDC-42 at basolateral contacts and thereby prevents recruitment of PAR-6 to these sites, at least two RhoGEFs, CGEF-1 and ECT-2, activate CDC-42, thereby counteracting PAC-1 (Chan and Nance, 2013). Whether and how these factors affect cortical flow at this stage and how other apical polarity determinants are regulated has not been analyzed so far. Moreover, *pac-1* loss-of-function embryos are viable despite transient mis-localization of PAR-6 (Anderson et al., 2008).

Previously, we demonstrated that shortly after the switch from anteroposterior to apicobasal polarization, an asymmetrically positioned midline forms in the *C. elegans* embryo through chiral morphogenesis, a rotational cell rearrangement with invariant directionality that is crucial for l/r symmetry breaking (Pohl and Bao, 2010). Chiral morphogenesis requires laterally asymmetric regulation of cortical contractility (Pohl and Bao, 2010) and is preceded by rotational cortical flows during division of the ectodermal blastomeres (Naganathan et al., 2014). The lateral asymmetry of cell movements and contacts strongly suggests that an unknown mechanism has to establish planar polarity at this stage. This elusive mechanism seems to utilize regulators involved in establishing apicobasal polarity, since chiral morphogenesis depends on CDC-42 (Pohl and Bao, 2010). Additionally, chiral morphogenesis also relies on non-canonical Wnt signaling (Pohl and Bao, 2010). The latter developmental signaling pathway has been shown to regulate embryonic spindle orientation (Schlesinger et al., 1999; Walston et al., 2004; Cabello et al., 2010; Sugioka and Bowerman, 2018) modulated by additional factors such as latrophilins (Langenhan et al., 2009) or syndecan (Dejima et al., 2015) and to determine cell fates in the early embryo (reviewed in Sawa and Korswagen, 2013) that seem to depend in part on Wnt-dependent induction of spindle asymmetry (Sugioka et al., 2011). Moreover, in *C. elegans* (Goldstein et al., 2006) as well as other systems (Habib et al., 2013), it has been shown that Wnt signaling can polarize isolated cells. In the *C. elegans* embryo, polarizing Wnt signaling emerges in the posterior blastomere, P1, and is then transduced from posterior cells to anterior cells by a relay mechanism that keeps re-orienting cells in the direction of the posterior polarizing center (Bischoff and Schnabel, 2006). This mechanism is most likely utilizing anteroposterior polarization of Wnt signaling components during mitosis as it has been shown that MOM-5/Frizzled is enriched at the posterior pole of cells before division in later embryogenesis (Park et al., 2004).

Hence, although Wnt signaling patterns the anteroposterior axis and is required for l/r symmetry breaking in *C. elegans*, roles of Wnt signaling in establishing planar cell polarity (PCP) have so far only been documented in neuronal morphogenesis. Here, Wnt signaling and the canonical PCP genes *fmi-1*/Flamingo, *prkl-1*/Prickle, *vang-1*/Van Gogh, *cdh-4*/Fat, *unc-44*/Diego regulate processes like fasciulation, neurite outgrowth, positioning and axon guidance (reviewed in Ackley, 2014). Importantly, neuro-morphogenesis as well as organogenesis in *C. elegans* results from interactions of individual cells with complex lineage trajectories and morphogenetic processes often occur in a local, piecemeal fashion (Bischoff and Schnabel, 2006; Rasmussen et al., 2008; Harrell and Goldstein, 2011; Pohl et al., 2012). Due to these developmental features, obvious planar polarized patterns based on polarity or PCP molecular markers have remained elusive for *C. elegans* embryogenesis.

In this study, we describe how planar polarized non-muscle myosin-aPAR domains form at the medial cortex at the time of l/r symmetry breaking in all the cells except P2. We find that non-muscle myosin-driven centripetal flow, which originates at cell-cell contacts, are at the core of this process. We characterize this process using particle image velocimetry (PIV), which reveals an anisotropy of flow emerging from anterior and posterior contacts. This in turn leads to the asymmetric positioning of non-muscle myosin-aPAR domains. We show that GSK-3 is essential for release of non-muscle myosin from cell-cell contacts, thereby directly affecting flow-dependent advection of aPARs. In one blastomere, ABpl, this leads to emergence of PCP through planar polarized localization of PAR-3 at a single cell-cell contact. For emergence of PCP, CDC-42 is required to regulate both PAR-3 and PAR-6 at cell-cell contacts, while PAC-1 is required for PAR-6 advection. We quantitatively describe additional molecular asymmetries at cell-cell contacts including cell adhesion and Wnt signaling components. This type of PCP is later re-deployed during pre-morphogenetic development in posterior lineages. Thus, a balance between contact retention and release of non-muscle myosin and aPARs seems to determine the degree of advection, thereby controlling planar polarization of cell-cell contacts and the apical domain, constituting the first instance of obvious PCP in the early *C. elegans* embryo.

## Results

### Advection and unmixing of apical cortical factors

*C. elegans* employs invariant l/r asymmetric contractility to establish an asymmetrically positioned midline through chiral morphogenesis (Pohl and Bao, 2010). This depends on the actin regulators Arp2/3 and CYK-1 (formin) as well as on non-canonical Wnt signaling (Pohl and Bao, 2010). Since previous studies imply roles of polarity factors in establishing apicobasal polarity at this stage (Nance et al., 2003; Anderson et al., 2008), we decided to explore how cortical contractile actomyosin flow impacts on them. We therefore quantitatively analyzed the dynamics of cortical and cell polarity factors (see Materials and Methods; Zonies et al., 2010; Dickinson et al., 2013; Heppert et al., 2018) with high-resolution time-lapse microscopy. We observed that – except for P2 – all cells at this stage show centripetally directed myosin flow, emanating from apical cell-cell contacts and collecting at the apical center to form a stable ring-like structure for ABpl (Figure 1A and B; Figure S1A and B; Video S1;) and crescent shaped structures for ABpr and MS (Figure S1C-F, Video S2). Centripetal flow seems equivalent to centripetal flow described during gastrulation (Pohl et al., 2012; Roh-Johnson et al., 2012). Concomitant with centripetal flow, PAR-6 accumulates at the center of the ring (Figure 1A, 200” to 300” time points; Figure 1B; Munro et al., 2004) and also adjacent to non-muscle myosin II, NMY-2, crescents (Figure S1C and D). Hence, PAR-6 forms a compact, transiently stable, apical domain unlike the previously reported uniform localization (Anderson et al., 2008).

**Figure 1.**
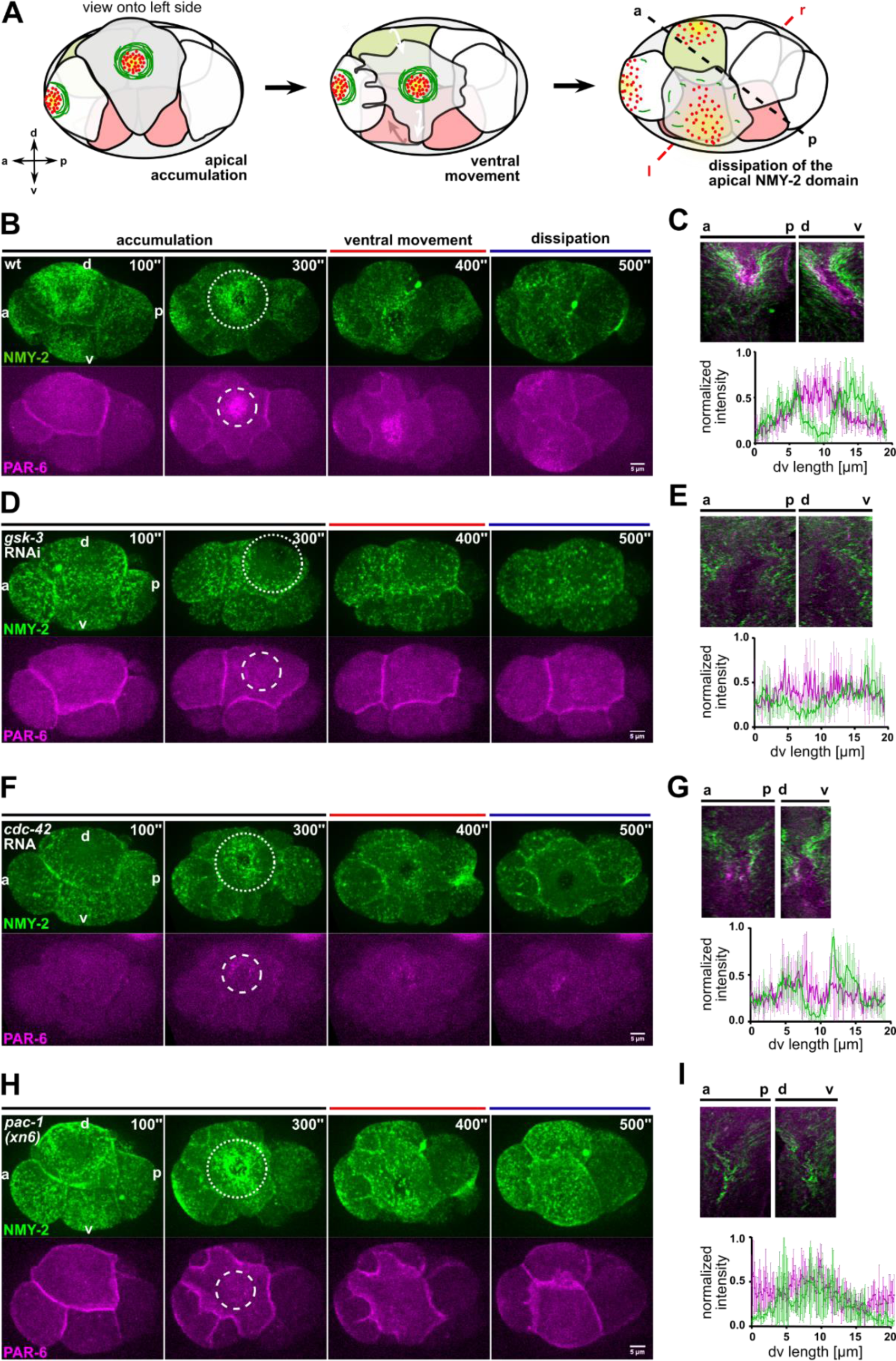
Centripetal actomyosin flow is essential for medial localization of PAR-6. **(A)** Schematic representation of the different stages of actomyosin flow on the left side of the embryo during chiral morphogenesis. Green depicts NMY-2, red PAR-6. **(B, D, F and H)** Representative time lapse images of projected apical sections of wt, *gsk-3* RNAi, *cdc-42* RNAi, and *pac-1(xn6)* embryos (shown is the left side) expressing NMY-2::GFP and mCherry::PAR-6 at the transition from 6-to 8-cell stage. Time is with respect to the completion of the ABp division. The axis directions are represented in the first timepoint. Dashed circles mark the NMY-2 and PAR-6 transiently stable apical structures, respectively. **(C, E, G and I)** Top: Kymographs of the apical cortex of ABpl along anteroposterior (a-p) and the dorsoventral (d-v) axis, showing the dynamics of cortical flow and localization of PAR-6 along time in wt (C), *gsk-3* RNAi (E), *cdc-42* RNAi (G) and *pac-1(xn6)* (I) animals. Bottom: Normalized intensity plots of NMY-2::GFP and PAR-6::mCherry along d-v axis of ABpl apical cortex in wt (C) (n = 5), *gsk-3* RNAi (E) (n = 3), *cdc-42* RNAi (G) (n = 4) and *pac-1(xn6)* (I) animals (n = 4).

Since cortical NMY-2 and PAR-6 dynamics are also influenced by cell cycle progression and the cell movements of chiral morphogenesis (Pohl and Bao, 2010), we decided to analyze this developmental stage by dividing it into three phases, (I) initial apical accumulation of NMY-2 (0-300 s after completion of ABp cytokinesis), (II) ventral movement of ABpl (300-400 s), which is influenced by EMS cytokinesis and (III) dissipation of apical NMY-2 and PAR-6, which is occurs during P2 division (400-500 s) (Figure 1A and B; Table S1).The accumulation of PAR-6 in the center of the NMY-2 ring occurs during centripetal flow (Figure 1C) and shows a time lag of 50±7 s (n = 4) relative to maximal accumulation of NMY-2 at the medial cortex (Figure 1B; Figure S1B). Based on previous modeling and experiments (Goehring et al., 2011; Mittasch et al., 2018) and since flow is fast enough (11±2 µm/min, n = 4) as well as aPARs associate with the cortex sufficiently long enough (PAR-6 recovery half-time is 7±2 s; n = 5), PAR-6 seems to be advected by centripetal cortical flow. Thus, while apical NMY-2 and aPARs accumulate together in the anterior half during polarization of the one-cell *C. elegans* embryo and stay segregated from PAR-2 only as long as flow persists (Cuenca et al., 2003; Munro et al., 2004), here, NMY-2 and PAR-6 unmix during centripetal flow and a medial PAR-6 domain persists for another 85±9 s (n = 4) after NMY-2 centripetal flow has stopped (Figure 1B and C).

### Regulation of centripetal cortical flow

Next, we asked whether centripetal flow is required for PAR-6 accumulation and performed a targeted screen (Figure S2) that included factors known to affect rotational flow and chiral morphogenesis (Pohl and Bao, 2010; Singh and Pohl, 2014; Naganathan et al., 2014). For one of these factors, GSK-3, we observed a strong loss of cortical flow and a complete lack of cortical NMY-2 ring formation (Figure 1D; Figure S3A; Video S3). Importantly, cell division timing in these *gsk-3* RNAi embryos is by and large normal for AB blastomeres and no cortical flow phenotype or fate switch has been found for the first two divisions in *gsk-3* RNAi (Fievet et al., 2013; Du et al., 2014), arguing that loss of centripetal flow after GSK-3 depletion is a specific phenotype at this developmental stage. Consistent with advection being responsible for PAR-6 accumulation, *gsk-3* RNAi embryos show a complete lack of PAR-6 apical accumulation and precocious dissipation (Figure 1D and E; Video S3).

Furthermore, we wanted to test the role of the Rho GTPase CDC-42, which has been implicated in the localization of polarity factors at the apical cortex (Anderson et al., 2008; Chan and Nance, 2013). For this, we performed partial depletion of CDC-42 by short-term RNAi (24-28 h). Under these conditions, NMY-2 puncta are significantly reduced (Figure 1F), though this does not abrogate centripetal flow of NMY-2 and a stable ring configuration is formed (Figure S3B; Video S4). However, as a consequence of reduced NMY-2 puncta and apparently reduced myosin flow, PAR-6 accumulation is also substantially reduced and, in most cases, no discernible domain is visible (Figure 1F; Figure S3B). Remarkably, lack of an apical PAR-6 domain leads to a collapse of the NMY-2 ring structure (in 22% of *cdc-42* RNAi embryos; n = 9), consistent with the idea that centripetal flow leads to a stable ring structure of NMY-2 due to unmixing and PAR-6 forming a stable, flow impenetrable domain after accumulation. These findings are consistent with previous findings of CDC-42 being essential for apical localization of PAR-6 at this stage (Anderson et al., 2008).

In addition, a role for the RhoGTPase activating protein (RhoGAP) PAC-1 in driving the switch from anteroposterior to apicobasal polarization has been described previously (Anderson et al., 2008). While *pac-1(xn6)* mutant embryos show normal centripetal NMY-2 flow (Figure 1H and 1I; Video S5), they completely lack advection of PAR-6 (Figure S3C). Hence, in accordance with the suggested role in apicobasal polarization (Chan and Nance, 2013), we find that *pac-1(xn6)* mutant embryos lack PAR-6 advection, however, they show higher cell-cell contact levels of PAR-6 (Figure 1H and see below). Similar to CDC-42 depletion, in 22% of *pac-1(xn6)* embryos (n = 9), the NMY-2 ring structure collapses (Video S5). Therefore, we conclude that while GSK-3 and CDC-42 are generally required for centripetal cortical flow at this stage, PAC-1 is specifically required for the release and advection of PAR-6 from cell-cell contacts, and both CDC-42 and PAC-1 are required to advect enough PAR-6 to generate a stable, flow-resistent medial apical PAR-6 cap.

In addition, we wanted to test the opposite situation, generally increasing myosin activity. We did so through *rga-3* RNAi, where NMY-2 assumes a network-like appearance that still forms an apical ring, however, where individual NMY-2 puncta are not discernible (Figure S4A; Video S6). In 20% of the cases (n = 10), the NMY-2 ring collapsed in spite of an accumulation of PAR-6 at the centre, arguing that the PAR-6 cap cannot resist exaggerated cortical flow. We further tested the role of ECT-2, a RhoGEF for CDC-42, which has been implicated in apicobasal polarization at this stage by activating CDC-42 at the cell cortex (Munro et al., 2004). Consistent with previous findings and similar to the phenotype observed for *cdc-42* RNAi, ECT-2 depletion results in the reduction of myosin puncta (Figure S4B; Video S7) and in a slight deregulation of cell-cell contact MLC-4 (Figure S4C and D), but a stable ring configuration is formed in 80% of the cases (n = 5). In line with the effect of CDC-42 depletion, *ect-2* RNAi also affects PAR-6 accumulation at the apical centre and the medial PAR-6 domain is highly reduced as compared to wild type (wt) (Figure S4B; Video S7). Thus, *ect-2* RNAi seems to partially phenocopy *pac-1(xn6)* phenotypes.

### Anisotropic centripetal flow generates planar polarized cortical domains

Intriguingly, although centripetal cortical flow seems to symmetrically emerge from cell-cell contacts to the apical center in all embryonic cells at first glance, NMY-2 dissipative structures show a distinct, cell-specific anisotropic pattern (Figure S1E and F; Video S2). To ascertain cell-type specific structures and to answer the question as to what molecular asymmetries give rise to cortical anisotropies, we decided to perform a detailed analysis of cortical flow in the cell driving this morphogenetic process, ABpl, through particle image velocimetry (PIV) (Figure 2A) (Dutta et al., 2015). Quantification of flow velocities revealed that centripetal cortical flow is in fact anisotropic with flow emerging from posterior and ventral contacts having higher velocities ≤ 16 µm/min while cortical flow emerging from anterior and dorsal contacts only reaches ≤ 6 µm/min (Figure 2B). If we translate these flow directions into angles (see wind rose plot description in Figure 2A), around 31% of the vectors are directing between 20-60°, exhibiting a bias towards the anterior direction (Figure 2B, red arc). PIV during ventral movement shows a strict ventral orientation of flow vectors which is also due to ventrally directed translocation of the cell itself (Figure 2B, middle). Subsequently, dissipation of cortical NMY-2 also occurs with a ventral oriented bias (Figure 2B, right).

**Figure 2.**
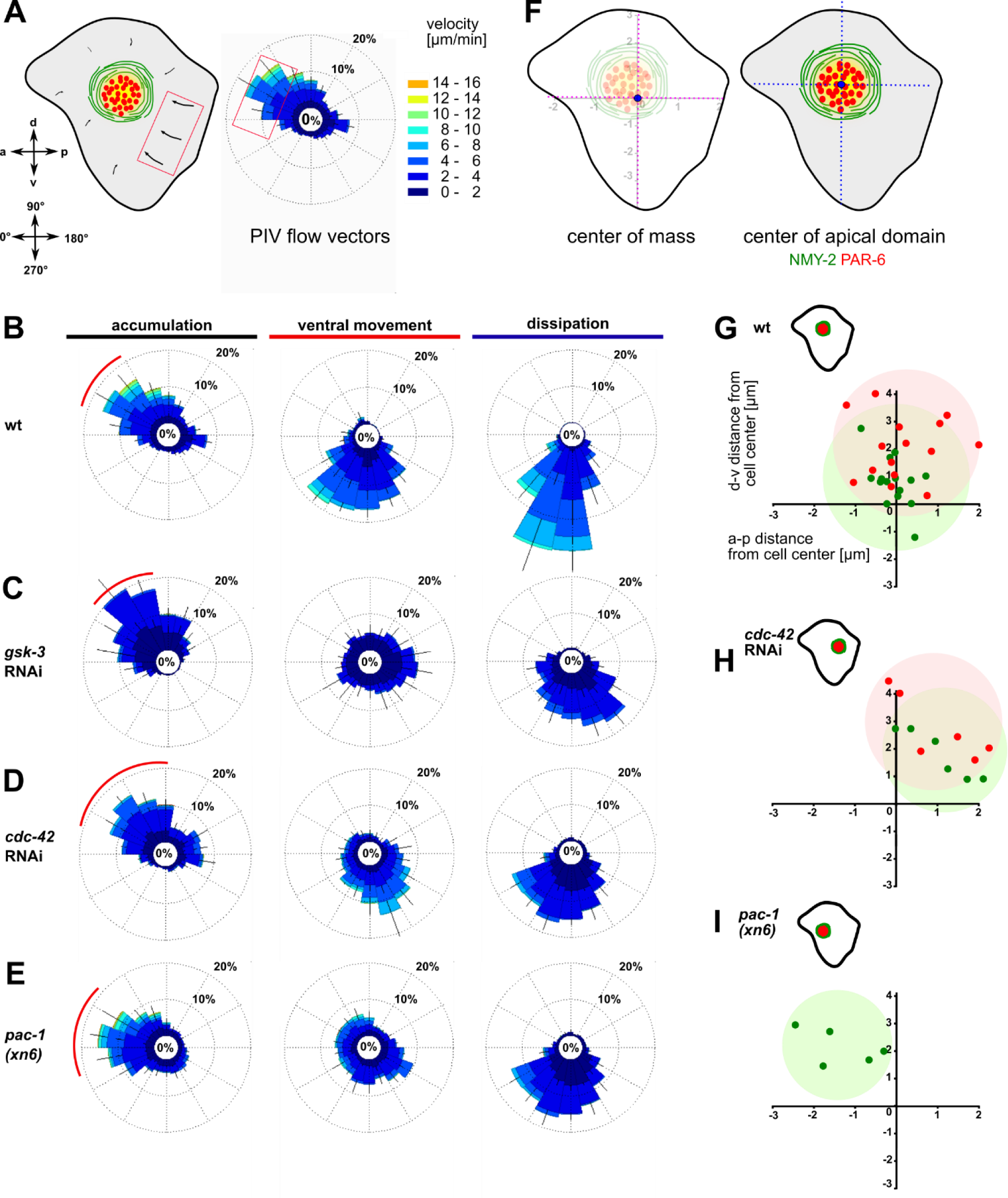
Anisotropic actomyosin flow generates planar polarized NMY-2-aPARs domains. **(A)** Left: Illustration of the ABpl apical cortex depicting the velocities of flow along a-p and d-v. Bottom: Axis direction correlated to angular coordinates. Middle: Wind rose plot depicting the direction and magnitude of cortical flow in the ABpl apical cortex generated by averaging velocities of NMY-2::GFP using PIV. Flow emerging from the posterior cell-cell contacts is represented in the anterior side of the plot and vice versa. Similarly, flow emerging from the ventral cell-cell contact is depicted in the dorsal side of the plot. Right: Color code for the magnitude of flow vectors. **(B-E)** Wind rose plots representing PIV analyses in wt (B), *gsk-3* RNAi (C), *cdc-42* RNAi (D) and *pac-1(xn6)* (E) animals (n = 3). The red arc depicts the bias in a specific direction during the accumulation phase of more than 30% of the vectors with high magnitude. **(F)** Left: Illustration of center of mass of ABpl apical cortex (blue circle). Right: Center of mass of NMY-2 (green circle) and PAR-6 (red circle) in the apical cortex with respect to the center of mass of the ABpl apical cortex (blue circle). **(G, H and I)** Insets: Illustrations of the positioning of the NMY-2-aPAR cortical domain of the ABpl cortex. Plots show the positioning of NMY-2 (green) and PAR-6 (red) in wt (G), *cdc-42* RNAi (H) and *pac-1(xn6)* animals (I); wt (n = 15) (G), *cdc-42* RNAi (n = 6) (H) and *pac-1(xn6)* (n = 5) (I); data points show center of the respective fluorescence signal with respect to the center of mass of the ABpl cell cortex which is taken as coordinate (0,0).

We next sought to ask whether factors responsible for the regulation of cortical flow also control its asymmetry. We analyzed cortical dynamics of embryos partially depleted for GSK-3, CDC-42, RGA-3 and ECT-2 as well as *pac-1* mutants. Consistent with a strong loss of centripetal flow in *gsk-3* RNAi embryos, PIV of cortical NMY-2 shows strongly reduced maximum velocities with almost 92% of the vectors lying in the ≤ 4 µm/min range and the anisotropy being shifted so that most of the flow vectors direct in the range of 40-80°, with a stronger dorsal bias (Figure 2C, red arc). A partial lack of polarity was also evident during the ventral movement and dissipation phase (Figure 2C, middle and right). Depletion of CDC-42 caused the flow to be reduced (≤ 12 µm/min) and the majority of flow vectors to be in the range of 20-90°, exhibiting a dorsal shift relative to wt (Figure 2D, red arc). In *pac-1(xn6)* embryos, the flow vectors as well as speeds increase in the 0-20° range, while the number of vectors and speeds decrease in the range of 40-60° (Figure 2E). Compared to wt having 25% of vectors with speeds of 4-6 µm/min in the range of 330-30°, flow vectors as well as speeds are increased in the 330-30° range with 44% having speeds of 8-10 µm/min, thereby exhibiting a strong anterior bias (Figure 2E, red arc).

We next asked if this anisotropy in NMY-2 flow affects the apical, cell type-specific dissipative structures and to this end closely looked at the positioning of the NMY-2-aPAR domain in ABpl (Figure 2F). We observed that apical domain formation in wt embryos had a slight bias towards the anterior side (Figure 2G), in agreement with the polarization of flow (Figure 2B, red arc). Further, we wanted to explore if any change in anisotropy could lead to changes in the positioning of the apical structure. To this end, we analyzed the factors in which cortical flow anisotropy was altered. Partial depletion of CDC-42 leads to reduced anterior flow velocities and a dorsal flow bias, causing a posterior shift of the domain (Figure 2H). In *pac-1(xn6)*, the domain is shifted anteriorly (Figure 2I), again in concert with findings that flow velocities now have a stronger anterior bias with higher velocities. Depletion of RGA-3 leads to a dorsal positioning of the domain relative to the wt due to exaggerated flow (Figure S4E) and partial depletion of ECT-2 leads to a posterior positioning of the domain similar to CDC-42 depletion (Figure S4F). These results suggest that both the magnitude and direction of cortical flow velocities might be playing key roles in determining medial apical position of the domain.

### Differential advection and contact retention of aPARs

Recent reports about the differential functions of the different aPARs made us look more closely at the localization and its possible functions (Rodriguez et al., 2017; Wang et al., 2017). To this end we asked whether all aPARs show the same distribution and dynamics as PAR-6. The use of endogenously tagged transgenes and live imaging pinpoints this differential regulation: In contrast to previous data that relied mostly on immunostaining (Anderson et al., 2008; Chan and Nance, 2013), live cell imaging shows a substantial population of endogenously tagged PAR-6 being constitutively localized to apical cell-cell contacts (Figure 1; Figure 3A; Figure S5A). Comparing PAR-6 to the aPAR kinase, the atypical iota type protein kinase C, PKC-3, and to the other aPAR PDZ domain protein, PAR-3, we found that while all aPARs are advected by centripetal flow, PAR-6 and PKC-3 are also partially retained at all apical cell-cell contacts of somatic blastomeres in the embryo (Figure 3A; Figure S5A; Video S8). In contrast, PAR-3 is only retained at a single cell-cell contact, the contact between ABpl and the P2 blastomere on the left side of the embryo (Figure 3A, bottom; Figure S5A, top; Video S8). PAR-3 is present at this contact shortly after completion of ABp cytokinesis and remains localized at this contact until the start of ventral movement of ABpl (Figure 3B, right; Video S8). Remarkably, although PKC-3 and PAR-6 are retained at all somatic cell-cell contacts at this stage, quantitative analysis reveals that they nevertheless show an anteroposterior polarity with an enrichment at the posterior cell-cell contact of ABpl (Figure 3B, left and middle). We speculate that posterior cell-cell contact localized aPARs must almost exclusively stem from the anterior cell (ABpl) since P2 only shows marginal levels of cortical aPARs much later when P2 divides into C and P3. Consistent with the idea that aPARs are advected by centripetal flow, we observe that PIV of PAR-3 in ABpl reveals the same directionality as NMY-2 flow, however, with slightly reduced velocities (Figure 3C). Considering the obvious and measured (Dickinson et al., 2017) differences between PAR-3 and PAR-6 cortical assemblies, it is interesting to note that we cannot detect a significant difference in the kinetics of advection to the medial cortex (Figure 3D).

**Figure 3.**
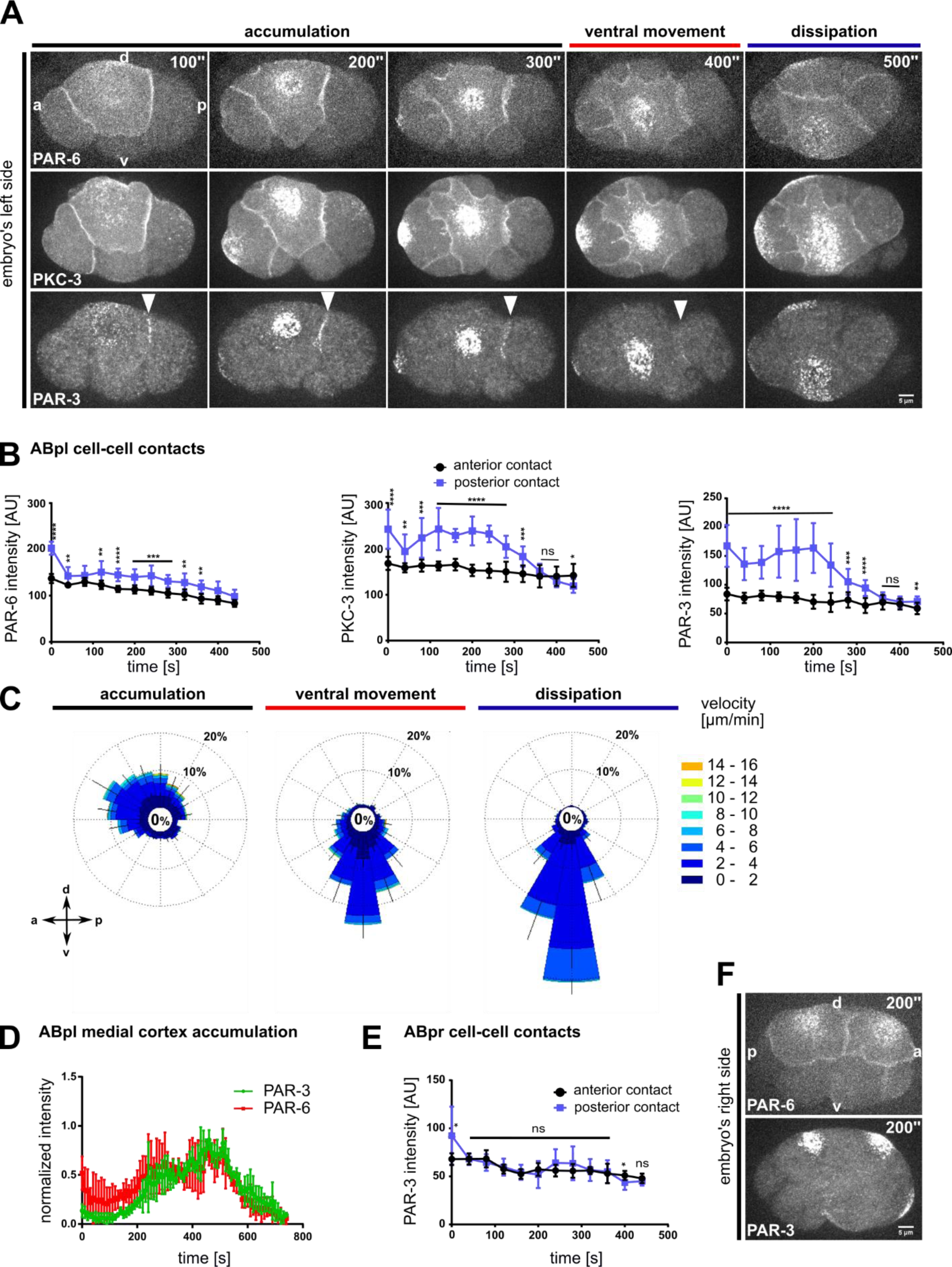
Advection and asymmetric contact retention of aPARs. **(A)** Representative time lapse images of apical cortical sections of embryos expressing mCherry::PAR-6 (top), GFP::PKC-3 (middle) and PAR-3::GFP (bottom). PAR-3 localization to the posterior contact (ABpl-P2) is marked by a white arrowhead. Time is with respect to the completion of the ABp division. **(B)** Quantification of mCherry::PAR-6 (n = 3), GFP::PKC-3 (n = 4) and PAR-3::GFP (n = 4) fluorescence intensities at the anterior (ABal-ABpl) and posterior (ABpl-P2) contact measured from apical cortical sections. Time is with respect to the completion of the ABp division. **(C)** Wind rose plots of PAR-3::GFP using PIV (n = 3). Right: Color code for the magnitude of flow vectors. **(D)** Normalized intensities of medial apical PAR-3 and PAR-6 fluorescence (n = 4). **(E)** Quantification of PAR-3::GFP fluorescence intensity at ABpr’s anterior (ABar-ABpr) and posterior (ABpr-P2) contact (n = 3). Time is with respect to the completion of ABp division. **(F)** Time lapse images of the right side of the embryo expressing mCherry::PAR-6 (top) and PAR-3::GFP (bottom). – All error bars indicate mean ± SD. P values: multiple t-test (*p < 0.05, **p < 0.01, ***p < 0.001, ****p < 0.0001).

Moreover, no obvious cell-cell contact localization of PAR-3 can be observed on the right side of the embryo, not in the fate-equivalent daughter of ABpl, ABpr (Figure 3E and 3F; Video S9). Hence, at the stage of l/r symmetry breaking, PAR-3 seems to be part of a unique mechanism of cell-cell contact regulation that is laterally specific and polarized according to the anteroposterior axis, therefore representing planar polarity.

### Regulation of aPAR cell-cell contact asymmetry

Next, we asked how regulators of centripetal cortical flow influence the cell-cell contact localization of NMY-2 and aPARs. We measured NMY-2 levels at the anterior and posterior cell-cell contacts of ABpl (the contacts ABal-ABpl, ABpl-P2, respectively) in wt and RNAi/mutant embryos (Figure 4A and B). Upon GSK-3 depletion, NMY-2 levels are increased at the contacts (Figure 4A, blue arrows) and non-muscle myosin regulatory light chain, MLC-4, levels are increased almost three-fold relative to wt levels (Figure S4C, middle). These data suggest that GSK-3 depletion leads to increased retention of the non-muscle myosin holo-complex, and, consequently, highly reduced apical flow. In contrast, CDC-42 depletion does not affect NMY-2 or MLC-4 levels at cell-cell contacts (Figure 4A and B; Figure S4C, right).

**Figure 4.**
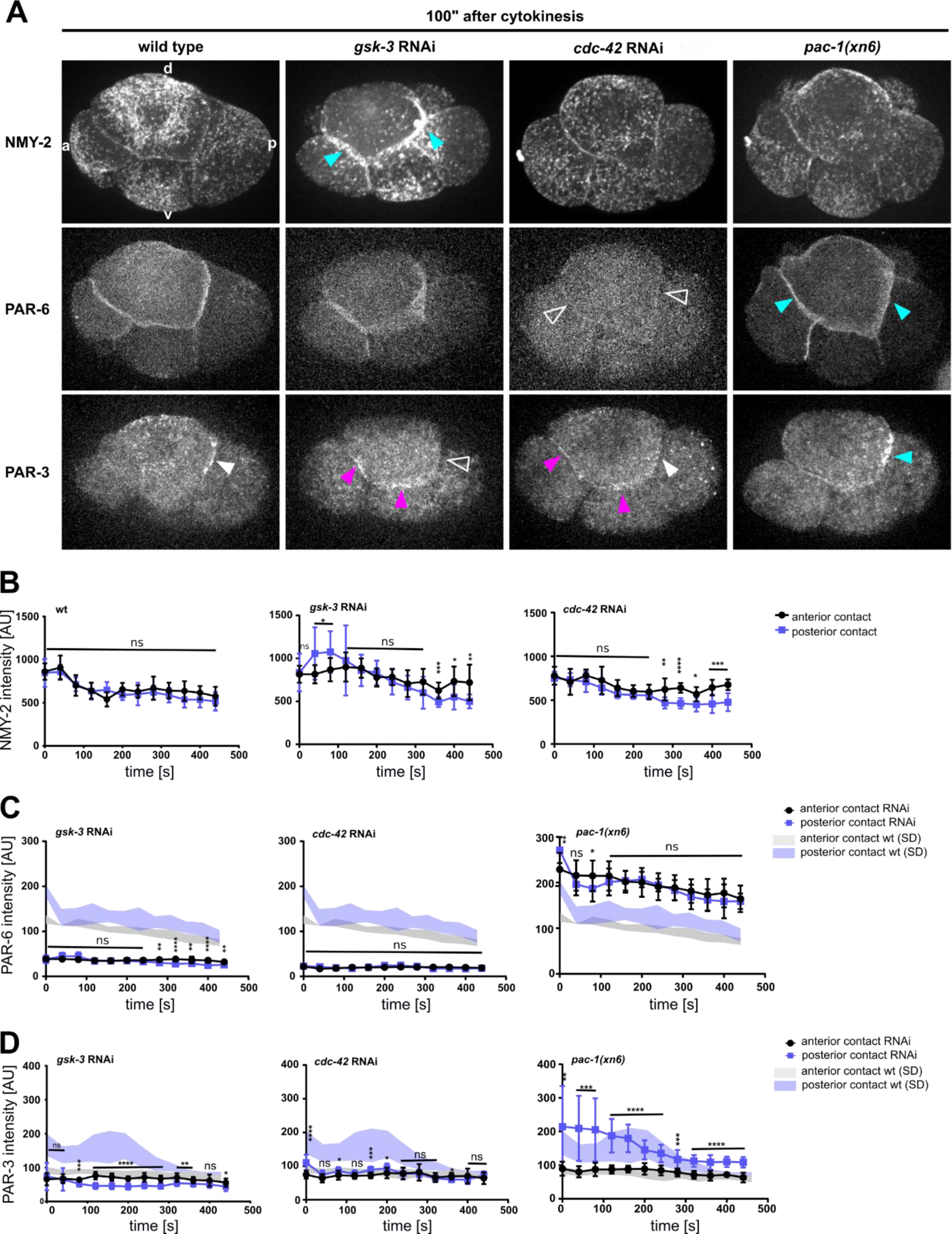
Regulation of NMY-2, PAR-6 and PAR-3 at the cell-cell contacts of ABpl. **(A)** Representative time lapse images of apical cortical sections of the left side of embryos expressing NMY-2::GFP (top), mCherry::PAR-6 (middle) and PAR-3::GFP (bottom) in wt and different RNAi/mutant backgrounds. Effects of the RNAi conditions on PAR-3 localization. White arrowhead: wt localization; empty arrowhead: loss of cell-cell contact localization; blue arrowhead: increased localization; fuchsia: ectopic localization. **(B-D)** Quantifications of NMY-2::GFP (B), mCherry::PAR-6 (C) and PAR-3::GFP (D) at ABpl’s anterior (ABal-ABpl) and posterior (ABpl-P2) contact measured from apical cortical sections of the indicated backgrounds (n ≥ 3). Time is with respect to the completion of ABp division. Error bars indicate mean ± SD. P values: multiple t-test (*p < 0.05, **p < 0.01, ***p < 0.001, ****p < 0.0001).

To ascertain factors essential for the unique cell-cell contact localization of PAR-3, we measured PAR-3 levels in RNAi/mutant backgrounds (Video S10). In contrast to wt embryos, where PAR-3 enrichment invariantly occurs at the posterior cell-cell contact of ABpl (Figure 3A; Figure 4A, arrowhead; Figure S5A), depletion of GSK-3 by RNAi leads to a loss of cell-cell contact-localized PAR-3 at the posterior side of ABpl and to a PAR-3 localization to the anterior and ventral cell-cell contacts (Figure 4A, bottom; Figure 4D; Figure S5B, top panels). In contrast, PAR-6 localization seems to be reduced significantly in all contacts (Figure 4A, middle; Figure 4C). Partial depletion of CDC-42 also leads to a loss of enrichment of PAR-3 at the posterior contact, thereby making the localization at both contacts more symmetric (Figure 4A, bottom, and 4D; Figure S5B). While CDC-42 seems to be very essential for PAR-6 as it becomes undetectable at cell-cell contacts in *cdc-42* RNAi (Figure 4A, middle; Figure 4C). Moreover, when quantifying medial accumulation of NMY-2, PAR-3 and PAR-6, we found that PAR-6 medial localization in *cdc-42* RNAi is highly reduced (Figure S5C). This is most likely due to lack of PAR-6 at cell-cell contacts already before initiation of centripetal cortical flow and consequentially less material to be advected in general (Figure 4A and C). However, medial localization of PAR-3 seems to be increased in the absence of CDC-42 (Figure S5B and D, grey bars). Our speculation is that PAR-3 retention at ABpl’s posterior contact is weakened upon CDC-42 depletion and more PAR-3 is available for advection. In accordance with this, in *pac-1(xn6)* animals, where cell-cell contact localized CDC-42 is rendered more active, this asymmetric PAR-3 localization is even increased (Figure 4A, bottom right; Figure 4D; Figure S5B), indicating higher retention (Anderson et al., 2008). Consequentially, medial localization of PAR-3 is decreased in *pac-1(xn6)*, pointing to less availability of PAR-3 at the apical cortex (Figure S5D, magenta bars). Thus GSK-3 and CDC-42 seem to be essential for cell-cell contact localization of PAR-3 and PAR-6. Moreover, the above data collectively suggest a coupling between contact retention and advection, which is partially defective in *pac-1(xn6)*. This coupling then brings about planar polarization of the apical cortex.

Previously, it was shown that two RhoGEFs, CGEF-1 and ECT-2 positively contribute to the cell-cell contact localization of PAR-6 after the switch from anteroposterior to apicobasal polarization by activating CDC-42 (Chan and Nance, 2013). Therefore, we performed partial depletion of ECT-2 by 8 h feeding RNAi and found reduced PAR-6 levels at cell-cell contacts (Figure S4D) and at the medial apical domain (Figure S5E). Furthermore, we found a properly established PAR-3 asymmetry, which is precociously lost (Figure S4D, middle). Hence, PAR-3 and PAR-6 have specific regulators and only partially share common regulators such as the RhoGEFs ECT-2 and potentially CGEF-1 that mediate retention of aPARs at cell-cell contacts.

### Cell-cell contact asymmetry of cortical regulators, signaling and cell adhesion molecules

The anisotropy in cortical flow (Figure 2), despite rather symmetric localization of NMY-2 and MLC-4 at cell-cell contacts (Figure 4B; Figure S4C), suggested that Rho GTPases and their regulators might be playing a major role in creating cortical and contact asymmetries. In this study as well previous studies, CDC-42 seems to be playing a major role in polarity establishment in conjunction with non-muscle myosin II and its regulators (Anderson et al., 2008; Chan and Nance, 2013). To determine the possibility that CDC-42 might be asymmetric at cell-cell contacts, we used a sensor for activated CDC-42, a GFP-tagged CRIB/G-protein binding domain of WSP-1 (Kumfer et al., 2010). We observed a significant difference in the distribution of active CDC-42 in the anterior and posterior contacts (Figure 5A), with higher levels on the posterior contact. This is consistent with a positive role of CDC-42 in cortical flow (Fievet et al., 2013), since we observe directional flow with the highest velocities from posterior (Figure 2B). Contrary to the previous reports (Anderson et al., 2008), however, we could not detect any cortically but solely cell-cell contact localized active CDC-42 (Figure S6).

**Figure 5.**
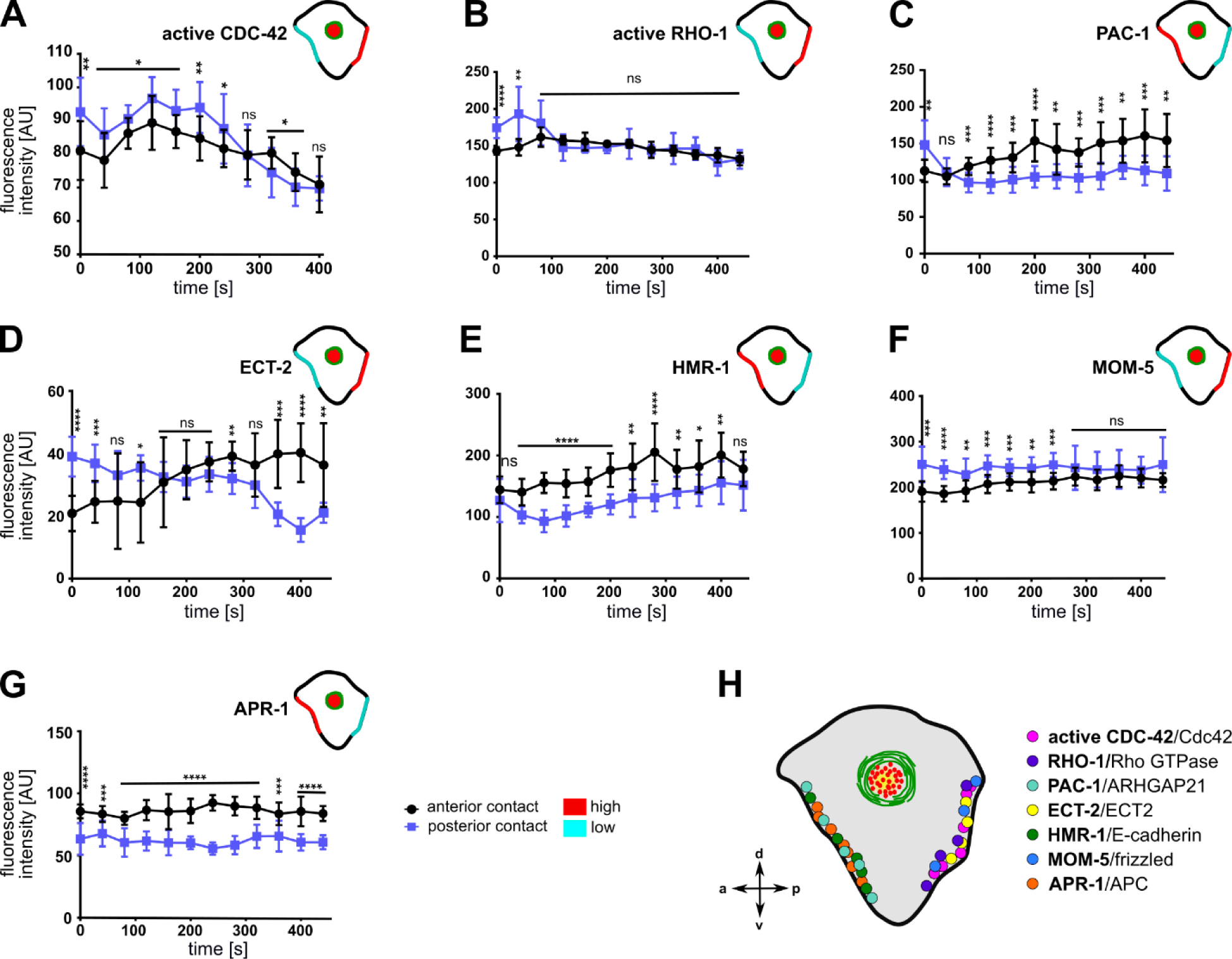
Asymmetric cell-cell contact localization of cortical regulators, signaling and cell adhesion molecules. **(A-G)** Quantification of fluorescence intensities of different transgenes at the anterior (ABar-ABpr) and posterior (ABpr-P2) contact of the ABpl apical cortex (n ≥ 3). Time is with respect to the completion of ABp division. Upper right inset: Illustration of the enrichment of the transgenes at the anterior and posterior contact; high: red, low: cyan. Error bars indicate mean ± SD. P values: multiple t-test (*p < 0.05, **p < 0.01, ***p < 0.001, ****p < 0.0001). **(H)** Illustration of the dominant localization of the different factors at both cell-cell contacts.

To examine the possibility that other Rho GTPases or their regulators might display an asymmetry at cell-cell contacts, we monitored a GFP-sensor for active RHO-1 (Figure 5B; Tse et al., 2012). It showed an initial posteriorly biased asymmetry but became symmetrically localized at cell-cell contacts in later timepoints (Figure 5B). Next, we sought to examine the cell-cell contact distribution of the RhoGAP PAC-1. We detected an enrichment of PAC-1 at the anterior contact compared to the posterior contact (Figure 5C). This is consistent with idea that PAC-1 curbs CDC-42 activity (Anderson et al., 2008; Chan and Nance, 2013) and with our data showing that CDC-42 is more active at the posterior cell-cell contact (Figure 5A). Moreover, quantifying a GFP-tagged ECT-2 transgene revealed an enrichment in the posterior contact compared to the anterior contact during accumulation phase, however, subsequently, asymmetry switches and ECT-2 is enriched in the anterior cell-cell contact (Figure 5D).

Since E-cadherins have been implicated in apicobasal polarization in several studies previously (Johnson et al., 1986; Nejsum and Nelson 2009; Stephenson et al., 2010; Klompstra et al., 2015), we also decided to examine its localization at the contacts. Upon quantification, we found a significant anterior enrichment of HMR-1, the E-cadherin ortholog (Figure 5E). This might relate to its role in translating specific contact cues into polarized recruitment of PAC-1 (Klompstra et al., 2015). Additionally, the Wnt pathway has been previously implicated in playing an essential role in chiral morphogenesis (Pohl and Bao, 2010) and MOM-5/Frizzled shows an anteroposterior asymmetric localization in cell divisions during later embryogenesis (Park et al., 2004). Therefore, we quantified MOM-5 levels at cell-cell contacts. Consistent with an anteroposterior polarization, we found that MOM-5 being asymmetrically localized at cell-cell contacts, with a posterior enrichment (Figure 5F). Furthermore, APR-1, the ortholog of APC in *C. elegans*, is enriched at the anterior cell-cell contact (Figure 5G) suggesting an opposing regulation of Wnt signaling at the anterior (Wnt low) versus posterior (Wnt high) cell-cell contact. These results are consistent with the anterior localization of APR-1 and posterior nuclear β-catenin localization in asymmetric seam cell divisions (Mizumoto and Sawa, 2007). Taken together, these results suggest that two opposing sets of protein complexes shape cell-cell contact asymmetry at this stage, one which is anteriorly and the other posteriorly enriched (Figure 5H).

### Morphogenetic role of planar polarized PAR-3 localization

The localization of PAR-3 to a single cell-cell contact during chiral morphogenesis prompted us to analyze PAR-3 localization during subsequent stages of embryogenesis. Intriguingly, we found that PAR-3 shows highly lineage-specific asymmetric cell-cell contact localization (Figure 6A-D). Specifically, the first contact that shows clear PAR-3 localization is the contact between ABp and P2 (Figure 6A). This localization is not restricted to one side of the embryo and thus also not planar polarized, however, it occurs prior to establishment of l/r asymmetry. As mentioned above, localization to the ABpl-P2 and later to the ABpl-C cell-cell contact is unique and not mirrored on the right side of the embryo by ABpr (Figure 6B). Subsequently, PAR-3 localizes to the MS-E cell-cell contact (Figure 6C; Video S11). This boundary between mesoderm and endoderm is maintained after the division of MS (Figure 6C). At the same developmental stage, on the other side of the embryo, PAR-3 localization between ABpl and C is lost during ABxx cell divisions but is rapidly re-established after completion of the divisions (Figure 6D). Slightly earlier, PAR-3 starts to accumulate at the C-P3 cell-cell contact. Like this, the C blastomere is almost completely encased by PAR-3-containing cell-cell contacts (Figure 6D; Video S12). Collectively, this shows that PAR-3 marks cell-cell contacts to generate a planar polarized pattern in the embryo that demarcates specific posterior lineages, E, C and P.

**Figure 6.**
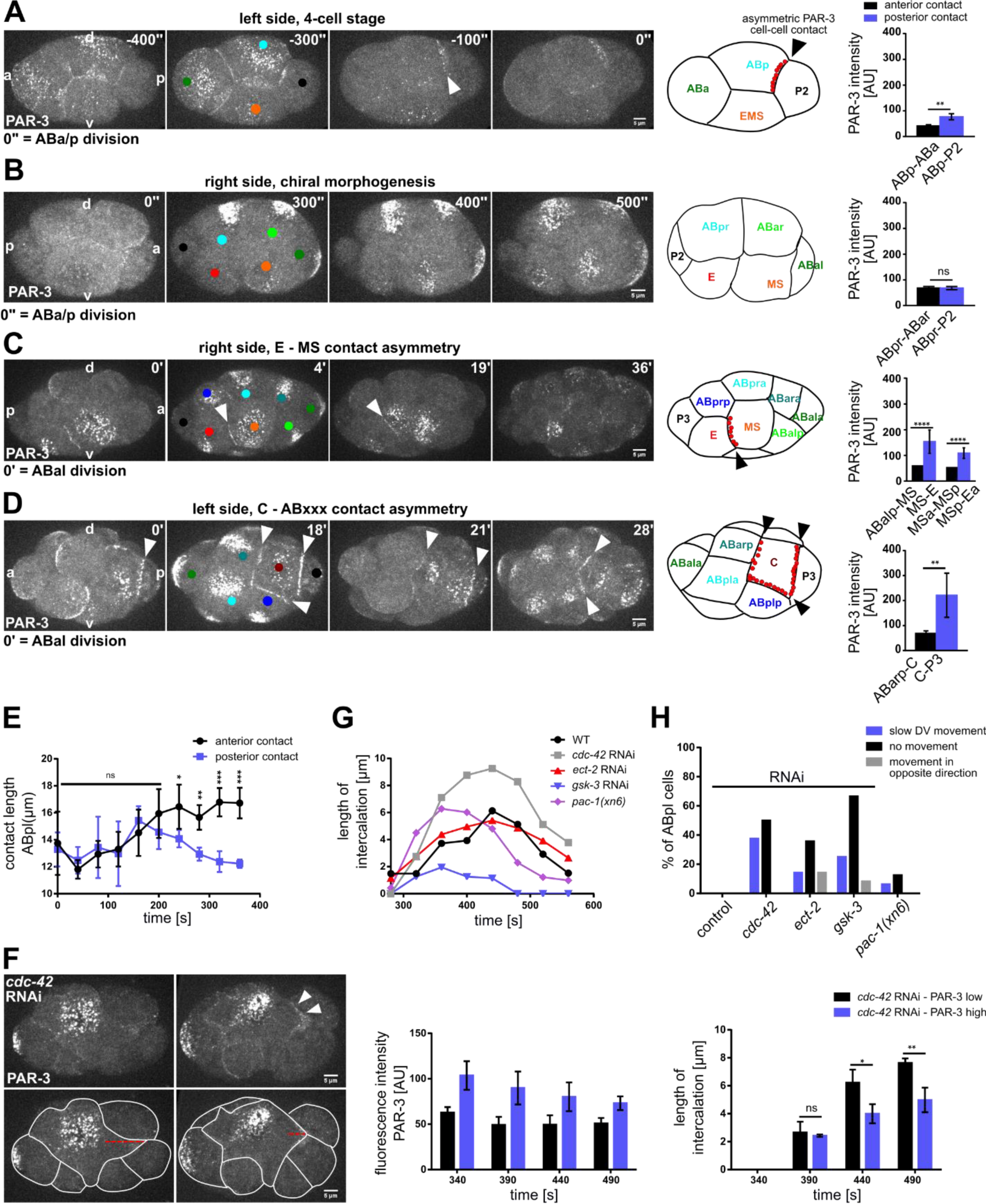
Morphogenetic role of cell-cell contact localized PAR-3 in posterior lineages. **(A-D)** Left: Representative time lapse images of cortical sections of embryos expressing PAR-3::GFP at 4 cell (A), 6-8 cell (B), 12-cell stage, right side (C), 12-cell stage, left side (D). White arrowheads point to cell-cell contact localized PAR-3. Cell identities are marked by colored circles, colors correspond to cell name colors in the embryo models (PAR-3 localization is marked in red). Right: Quantification of PAR-3::GFP fluorescence intensity at anterior and posterior contacts of ABp (A) (n = 2), ABpr (B), MS and MSp (C) (n = 3), C (D) (n = 3), measured from apical sections. **(E)** Length of the anterior (ABa-ABp) and posterior contact (ABp-P2) at the ABpl apical cortex (n = 4). **(F)** Top left: Representative time lapse images of cortical sections of *cdc-42* RNAi embryos expressing PAR-3::GFP either at low level (upper left) or high level (upper right) at the posterior contact. Bottom left: Intercalation lengths of ABpl during the P2 division for both the above mentioned conditions. Middle: Quantifications of PAR-3::GFP fluorescence intensity in *cdc-42* RNAi embryos with low PAR-3 levels (black) and high PAR-3 levels (violet) at the posterior cell-cell contact. Right: Length of the intercalation of ABpl into the P2 furrow for the embryos quantified in the middle panel (n = 3). Time is with respect to the completion of ABp division. Error bars indicate mean ± SD. P values: multiple t-test (p < 0.05, **p < 0.01, ***p < 0.001, ****p < 0.0001). **(G)** Changes of the length of the intercalation of ABpl during the P2 division for different RNAi and mutant conditions over time (n = 4). Time is with respect to the completion of ABp division. **(H)** Quantification of different effects of the RNAi/mutant conditions on the ventral movement of ABpl during chiral morphogenesis (n ≥ 12).

**Figure 7.**
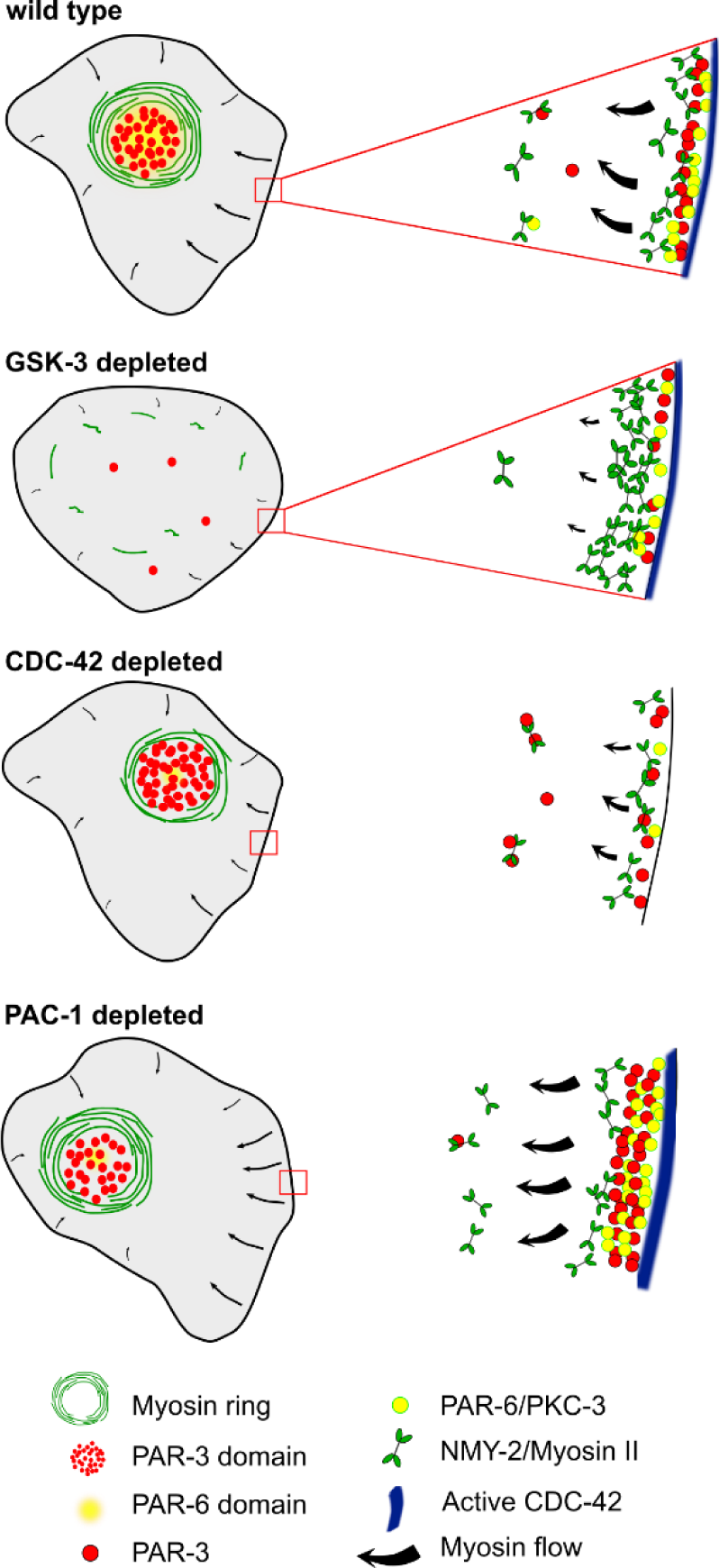
A model for the emergence of planar polarized myosin-aPARs medial domains. See discussion for details.

To better understand the roles for PAR-3 contact localization, we decided to study the ABpl-P2 cell-cell contact in more detail. Unlike the other blastomeres in the embryo at this stage, the cell-cell contacts of ABpl undergo a highly asymmetric development during the ventral movement of the cell, the anterior contact expands while the posterior shrinks (Figure 6E). We therefore asked how PAR-3 localization affects contact dynamics. When comparing wt and *cdc-42* RNAi embryos, we found that the contact between ABpl and P2/C is shows very different dynamics (Figure 6F, left). Unlike wt embryos, ABpl fully intercalates into the furrow of the dividing P2 cell in *cdc-42* RNAi embryos (Figure 6G; Video S4). Due to the variability in depletion by RNAi, we analyzed *cdc-42* RNAi embryos with strongly reduced ABpl-P2 cell-cell contact localized PAR-3 and compared these to *cdc-42* RNAi embryos with moderate reduction of PAR-3 (Figure 6F, middle). Consistent with the idea that PAR-3 contributes to separation of anterior from posterior cells, we found that embryos with strong PAR-3 reduction also show significantly increased erroneous furrow intercalation (Figure 6F, right). When averaging over several embryos, *cdc-42* RNAi embryos show a substantial increase in ABpl-P2 furrow contact length (Figure 6G) while this is less pronounced in *ect-2* RNAi embryos and almost not changed in *pac-1(xn6)* embryos (Figure 6G). We suggest that this is due to decreasing impact of these backgrounds on asymmetric PAR-3 cell-cell contact localization in the order *cdc-42, ect-2, pac-1* RNAi (Figure S5B). Previously, using laser irradiation-inflicted cell cycle delays, we could show that intercalation of ABpl into the EMS furrow is essential for proper movement of ABpl and generation of a tilted midline (Pohl and Bao, 2010). Consistently, ABpl fails to move ventrally when it retracts from the EMS furrow in all *gsk-3* RNAi embryos (Video S3) as well as *ect-2* RNAi embryos (9/14). Taken together, the level of erroneous furrow intercalation seems to be proportional to the number of embryos where chiral morphogenesis fails (Figure 6H): in *cdc-42* and *ect-2* RNAi embryos mostly due to slowing down of ventral movement by erroneous intercalation of ABpl into the P2 furrow, and in *gsk-3* and *ect-2* RNAi embryos due to complete lack or instability, respectively, of intercalation of ABpl into the EMS furrow (Figure 6H).

Finally, based on these findings, we decided to investigate whether aside from regulating cytokinetic cell intercalation at the ABpl-P2/C contact, cell-cell contact localized PAR-3 also regulates cytokinetic cell intercalation during later stages. To do so, we focused on the MS-E contact that also shows a very clear PAR-3 enrichment (Figure 6C). Here, during cytokinesis of the MS blastomere, neighboring AB-derived blastomeres readily intercalate into the MS furrow while E does not (Figure S7, white outlines). Moreover, we also observe that a large fraction of cortical PAR-3 accumulates in the midbody in the subsequent division of the E blastomere (Figure S7; arrowhead) and the remaining apical PAR-3 stays associated with the Ea-Ep cell-cell contact. This is only observed for the E blastomere and not for any other blastomere, arguing that clearing of apical PAR-3 through this mechanism might help to prepare Ea and Ep for gastrulation where they are covered by neighboring cells (Nance et al., 2003). Thus, a main function of cell-cell-cell contact localized PAR-3 seems to regulate cytokinetic cell-cell intercalation which can give rise to substantial cell rearrangements in a structure that is only composed of a few cells.

## Discussion

### Early establishment of planar polarity in *C. elegans*

More than 20 years ago, the concept was established that blastomeres in *C. elegans* are specified by a process of stepwise, binary diversification involving the Wnt pathway genes *lit-1*/NLK and *pop-1*/TCF/LEF (Kaletta et al., 1997; Lin et al., 1998). Subsequently, it was shown that the core of this specification system is a relay of Wnt-dependent spindle-polarizing information that originates in the germline blastomere P1 and is maintained in its descendants (Bischoff and Schnabel, 2006). Since the *C. elegans* embryo can be considered a squamous-like epithelium, this specification system will most likely require planar polarized domains to prevent cell-cell mixing at cell fate boundaries and during cell division, particularly since cell divisions usually generate anteroposteriorly staggered configurations. Here, we demonstrate that after the transition from anteroposteriorly polarized blastomeres in the 2-cell to apicobasally polarized blastomeres in the 4-cell embryo, specific blastomeres also become polarized in the plane of the embryonic epithelium. We show that planar cell polarity is established through deployment of the same machinery that patterns anteroposterior and apicobasal polarity, cortical contractile actomyosin flow together with anterior/apical polarity determinants, most importantly, PAR-3.

In addition to the first planar polarized cell, ABpl, which drives l/r axis formation, planar polarization of cells is restricted to cell contacts with posterior lineages that need to give rise to (mostly) clonal tissues, germline (P2 and P3), endoderm (E and Ex) and laterally symmetric body muscle (C and Cx). We present evidence that planar polarized landmarks on these lineages seem to help in preventing cell-cell intercalation during division of these lineages (Figure 6; Figure S7). In many embryos, regulated, cell division-mediated intercalations contribute to cell movements and patterning during early development, including the Drosophila (Founounou et al., 2013; Herszterg et al., 2013; Guillot and Lecuit, 2013), the chick (Firmino et al., 2016), and the Xenopus embryo (Hatte et al., 2014). This is due to the fact that cytokinesis has to adapt to the multicellular context, where the dividing cell biomechanically signals the need for adhesion remodeling to the neighboring cells (Herszterg et al., 2014). The situation in *C. elegans* is slightly different than in those organisms since furrowing is asymmetric and progresses from apical to basal, where the midbody is then localized, while in many other organisms, midbodies of embryonic epithelia end up on the apical side (Herszterg et al., 2014). This difference is most likely due (1) to the lack of polarized apical junctions in *C. elegans* that can serve as a mount for the actin cytoskeleton in other organisms, and (2) since the early *C. elegans* embryo is topologically different from other embryonic epithelia, consisting of a small number of squamous-like blastomeres where cell-cell contact rearrangements appear more similar to those in early embryos of other holoblastically cleaving species like mouse or human. Interestingly, although a stochastic process, lineage segregation depends on the inheritance of the apical domain in the mouse embryo (Maître et al., 2016; Korotkevich et al., 2017), highlighting a conserved function of apical polarity determinants in cell fate specification.

Our data support parts of our earlier model on the integration of mechanisms leading to a continuum of axial patterning in *C. elegans* (Pohl, 2015): It was previously shown that Wnt signaling, known to regulate chiral morphogenesis, is also required for chiral, counter-rotating flow during skewing of the ABa/ABp division (Naganathan et al., 2014). Based on these findings, we propose that directional cortical flow during cytokinesis of ABp might bias the distribution of Wnt pathway components such as MOM-5/Frizzled to become enriched on the ABpl/P2 interface (Figure 5F). This in turn might lead to anterior enrichment of antagonistically acting factors like APR-1/APC (Figure 5G; Mizumoto and Sawa, 2007). It seems plausible to speculate that factors acting downstream on cortical flow and aPAR advection/retention such as CDC-42 and its regulatory GAPs and GEFs receive instructive inputs from asymmetrically localized Wnt signaling as Wnt signaling has been shown to polarize other cytoskeletal structures such as the spindle (Goldstein et al., 2006; Sugioka et al., 2011; Sugioka et al., 2018). Thus, the intrinsic chirality of actomyosin dynamics during cytokinesis together with the impact of polarized Wnt signaling might constitute the main driver of axial patterning coordination once cell-cell contacts exist in the embryo (Sugioka and Bowerman, 2018).

### Role of Rho GTPases and their regulators in PCP

Previously, a role of cortical flow in controlling clustering of aPARs has been described for the polarization of the anteroposterior axis (Wang et al., 2017). Here, cortical flow enables clustering of PAR-3 as a response to cortical actomyosin contractility-generated tension. Moreover, reduced activity of CDC-42 allows the other aPARs, PAR-6 and PKC-3, to associate with PAR-3 clusters, while increased CDC-42 activity leads to a more diffuse cortical localization of PAR-3 and dissociation of aPAR co-clusters (Wang et al., 2017). Vice versa, PAR-3 clustering has been shown to be required for effective advection (Dickinson et al., 2017). Moreover, consistent with CDC-42 activity shaping aPAR complexes, formation of clustered versus diffuse aPAR complexes during anteroposterior polarization also depends on an inverse activity state of PKC-3 (Rodriguez et al., 2017), giving rise to clustered PAR-3-PAR-6-PKC-3^inactive^ (corresponding to the co-clustered aPAR complex with CDC-42^low^; Wang et al., 2017) and diffuse CDC-42-PAR-6-PKC-3^active^ (corresponding to aPAR co-cluster dissociation or CDC-42^high^, Wang et al., 2017). Although a different developmental stage, our data strongly support this type of aPAR complex regulation: In the first cell with planar polarized PAR-3 at cell-cell contacts, ABpl, we find that CDC-42 activity is presumably high in the posterior cell-cell contact due to the CDC-42-inactivating GAP, PAC-1, showing the reciprocal planar polarity of PAR-3 (Figure 5A and C). Notably, also active RHO-1 is initially enriched posteriorly (Figure 5B). Consistent with the findings during anteroposterior polarization, this would lead to dissociation of aPAR co-clusters at the posterior cell-cell contact, where active CDC-42 is localized (Figure 5A). This also explains, why not only PAR-3 but also PAR-6 and PKC-3 show a planar polarized localization, although not as pronounced as PAR-3 (Figure 3B). Accordingly, we find that PAR-3 is more readily advected and lost from ABpl’s posterior contact when CDC-42 levels are down-regulated (Figure S5D). Therefore, it seems plausible that when cortical flow emerges in vicinity of cell-cell contacts (where CDC-42 is no longer detectable), centripetal cortical flow might again trigger aPAR co-clusters that we find to be co-advected to the medial cortex (Figure 3D). However, unlike during the anteroposterior polarization, centripetal cortical flow is not able to advect all PAR-6 and PKC-3 from cell-cell contacts, which can be attributed to the interaction with contact-localized CDC-42 and interaction with cell-cell adhesion complexes that obviously did not exist in the one-cell stage. Remarkably, PAR-6’s interactions with cell-cell contact-localized factors seems to be specifically regulated by PAC-1, which, when mutated leads to loss of PAR-6 advection by centripetal flow, also from contacts with lower levels of PAC-1 (Figure 1H). These findings are mostly consistent with previous data (Klompstra et al., 2015), showing a multi-component protein complex scaffolded by E-cadherin recruiting PAC-1 to cell-cell contacts. Interestingly, we find that ABpl’s anterior cell-cell contact shows significantly higher HMR-1/E-cadherin levels than the posterior, which can explain the observed anterior PAC-1 enrichment (Figure 5E). We can only speculate that this asymmetric localization of PAC-1 might also contribute to enhanced levels of cell-cell contact F-actin and reinforce recruitment of cell-cell adhesion proteins as described for late stages of embryonic morphogenesis (Zilberman et al., 2017).

### Similarity and difference to other forms of PCP

During gastrulation in Drosophila, correct anteroposterior patterning of the extending germband requires a planar polarized pattern of non-muscle myosin II localizing to anteroposterior cell-cell contacts while Bazooka/PAR-3 localizes to dorsoventral contacts (Zallen and Wieschaus, 2004). Although PAR-3 localizes to the posterior cell-cell contacts in ABpl, the lack of hexagonal epithelia that are mostly controlled by junction mechanics-dependent neighbor exchanges, makes it difficult to compare the role of planar polarized cell-cell contacts in *C. elegans* to those in the early fly embryo. However, molecularly, there seem to be several similarities. For instance, it has been shown that for sensory organ precursor cells (SOPs) in the notum epithelium, PCP depends on the canonical PCP pathway involving, among others, *fz/frizzled, dsh/disheveled, Vang/Van Gogh*, and *fmi/Flamingo* (reviewed in Yang and Mlodzik, 2015). In the absence of PCP, SOPs divide with properly segregated antagonistic polarity domains (aPARs versus Pins/Numb), however, randomly with respect to the epithelial plane (Besson et al., 2015). Interestingly, aPAR domains already polarize before mitosis in dependence on Wnt/PCP. Therefore, similar to our data on the emergence of planar asymmetries of aPARs in the early *C. elegans* embryo, there also seem to be Wnt/PCP-dependent mechanisms that operate outside of their known roles in mitosis and spindle orientation (Yang and Mlodzik, 2015). Moreover, in the Drosophila ommatidial epithelium, PCP controls the unilateral localization of Bazooka/PAR-3, independently of Par-6 (Aigouy and Le Bivic, 2017), again highly similar to the pronounced asymmetry of PAR-3 at posterior cell-cell contacts in *C. elegans* that does not in all cases require proper regulation of PAR-6, for instance in *pac-1(xn6)* (Figure 4A).

In vertebrates, Par3’s role in planar polarity has been reported to be either uncoupled from or coupled to its role in apicobasal polarity, depending on the context. During mouse inner ear development, Par3 is asymmetrically localized in dependence on canonical PCP and Rac signaling but independently of Par6 and aPKC, moreover, it does not control spindle positioning through LGN/Gαi (Landin Malt et al., 2019). Interestingly, it has also been recently shown that Par3 might have an instructive role in PCP by direct binding to the core canonical PCP component Prickle3 during establishment of PCP in the Xenopus neural plate (Chuykin et al., 2018). On the other hand, Par3-dependent apicobasal polarity seems to be required to set up PCP in avian embryos (Lin and Yue, 2018). Thus, Par3/PAR-3 seems to constitute an evolutionarily conserved, context-dependent driver of PCP, either by establishing biomechanical planar polarity, relaying apicobasal polarity to planar polarity, reinforcing canonical PCP signaling, or helping to establish asymmetric localization of PCP components. Our data reveal that the early *C. elegans* embryo also requires PAR-3-dependent PCP to achieve proper signal integration and relay during axial patterning.

## Materials and methods

### *C. elegans* strain maintenance

Strains were maintained on standard Nematode Growth Media (NGM) as previously described (Brenner, 1974) and were cultured at 20-25°C. Strain names and genotypes used in this study can be found in Table S2.

### Mounting and dissection of embryos

Embryos were dissected from gravid hermaphrodites in M9 buffer on a cover slide. Embryos were selected at the 4-cell stage and mounted on a #1 coverslip (Corning, Lowell, MA) along with a 1 µl suspension containing M9 and 20 µm diameter poly-styrene microspheres (Polyscience, Warrington, PA). The preparation was then covered with another coverslip and sealed using vaseline. This sample preparation was imaged under the microscope usually starting from the ABa/ABp divisions.

### Live cell imaging

Appropriately staged embryos were imaged using a VisiScope spinning disk confocal microscope system (Visitron Systems, Puchheim, Germany) consisting of a Leica DMI6000B inverted microscope, a Yokogawa CSU X1 scan head, and a Hamamatsu ImagEM EM-CCD. Z-sectioning was performed with a Piezo-driven motorized stage (Applied Scientific Instrumentation, Eugene, OR). All acquisitions were performed at 20-23°C using a Leica HC PL APO 63X/1.4-0.6 oil objective. For imaging/quantifying of anterior and posterior contacts of ABpl, we collected z-sections of 16 focal planes (0.5 µm apart) with 10 s intervals with a 488 and 561 nm laser at an exposure of 150 ms from the onset of ABa/ABp division, for a total duration of 10 min. While for PIV analysis, we collected 8 z-sections (0.5 µm apart) with 3 s intervals, again for a total duration of 10 min with the same laser settings as above. For imaging 12-cell stage embryos, we collected 26 z-sections (1µm apart) with intervals of 1 min. For most imaging, laser intensities used were 5% for the 488 nm and 40% or 60% for the 561 nm laser (each 25 mW maximal output at the source).

### RNA interference

RNAi experiments were performed by feeding as previously described (Kamath et al., 2001) with a few modifications in the amount of time that the animal is kept on the plate according to the lethality of the gene targeted (see below). RNAi feeding bacteria were grown overnight (around 16-18 h) in 1 ml Luria broth with ampicillin at a concentration of 100 µg/ml and 500 µl of this culture was used to inoculate 10ml of LB ampicillin and grown at 37°C for 6-8 h. This culture was then centrifuged and resuspended in 300 µl of the same media, which was plated and kept for drying and induction on feeding plates (NGM agar containing 1 mM IPTG and 100 µg/ml ampicillin). Worms were kept on these feeding plates for the number of hours specified below and then dissected and mounted for imaging.

Weak RNAi perturbations of essential genes were performed by lowering the number of hours of feeding. The RNAi clone for *gsk-3* was availed from the Vidal library (Rual et al., 2004) and early L4 worms were kept on feeding plates for 24-36 h at 21-23°C and this was the temperature used throughout unless specified otherwise. *cdc-42, rga-3, ect-2* feeding clones were also obtained from the Vidal library. For *cdc-42* RNAi, L4 or young adults were kept on feeding plates for 23-25 h. For *rga-3* RNAi, young adults or adults were kept on feeding plates and 8-12 h of feeding was enough to show phenotypes and longer hours resulted in severe phenotypes. For *ect-2* RNAi, adults were kept on feeding plates for 12 h.

### Flow velocity analysis using PIV

We used PIV to track NMY-2 and PAR-3 particles/foci in the apical cell cortex of ABpl and to measure velocity distributions with high spatial resolution. The imaging conditions we used for the PIV analysis were with high temporal resolution of 3 s intervals and with a high axial resolution of 0.5 µm spanning 8 z-stacks, enough to span the entire apical cortical section of ABpl. These z-stacks were projected using ImageJ and the image series was loaded into the PIVlab MATLAB algorithm (Thielicke and Stamhuis, 2014; pivlab.blogspot.de). In brief, a grid is drawn on each image. A fixed size window centered at each grid-point defines the region of interrogation. Fast Fourier transform is used to calculate the cross-correlations of this region with regions in the subsequent image. The PIV analysis was performed by using a 3-step multi pass, 64X64 pixel (9.152 × 9.152 µm), 32 × 32 pixel (4.576 × 4.576 µm) and the final interrogation window of 16 × 16 pixels with 50% overlap. Only the area within the apical cortex boundary of ABpl was taken for the analysis. The vector profiles generated using PIV gave us information about the direction and magnitude of the particle/foci movements. The vector fields across time were generated from each stage of a single embryo – accumulation, ventral movement and dissipation. The time-averaged vector fields from each stage were averaged for 3-4 embryos and are represented by wind rose plots.

### Measurements

#### Fluorescence intensities and data analysis

All quantifications of fluorescence intensities of proteins were performed on maximum intensity projections of apical cortical sections. For all measurements, background intensities were subtracted from the integrated intensity of the signals. For all measurements of fluorescence intensities at cell-cell contacts, the imaging conditions are mentioned above. We quantified the integrated intensity in circular ROIs of 0.327 µm at the cell-cell contacts of ABpl/ABpr using ImageJ. This diameter was small enough to span the entire width of the contact. For a single time point, we quantified 3 ROIs along the contact. The same ROI was used to measure the background intensity, which was subtracted from the signal intensity. For most of the measurements of cell-cell contact fluorescence intensities, we quantified 4 embryos. For medial cortex intensity measurements, we again quantified 3 circular ROIs of 0.327 µm for integrated intensity at every time point indicated. Most of the statistical analysis was performed using multiple t-test.

#### Center of mass calculation

We calculated the center of mass of the apical cortex of ABpl cell and the NMY-2 and PAR-6 domains using an ImageJ plugin. We manually traced the apical cortex boundary of the ABpl cell from z-projected stills to calculate the center of the mass of the apical cortex. We also manually traced the NMY-2 and PAR-6 domains similarly. We then subtracted the ABpl cell center of mass coordinates from the coordinates derived from center of mass of NMY-2 and PAR-6 domains to give us the center of mass of the domains relative to the center of mass of the cell.

#### Intercalation lengths

We measured the intercalation lengths by measuring the distance from the apex of the intercalating lamellipodium to the edge of the lamellipodium from which the ABpl cell body starts.

## Acknowledgements

We acknowledge funding by the Cluster of Excellence Macromolecular Complexes in Action (Deutsche Forschungsgemeinschaft project EXC 115) and the LOEWE Research Cluster Ubiquitin Networks to C.P. We would like to thank all members of the Pohl Group for their support and helpful discussions. We would like to thank Michael Glotzer for providing the MG644 strain. Most of the strains used in this study were provided by the Caenorhabditis Genetics Center, which is funded by the National Institutes of Health Office of Research Infrastructure Programs (P40 OD010440).

## Supplementary material

### Supplementary figures

**Figure S1.**
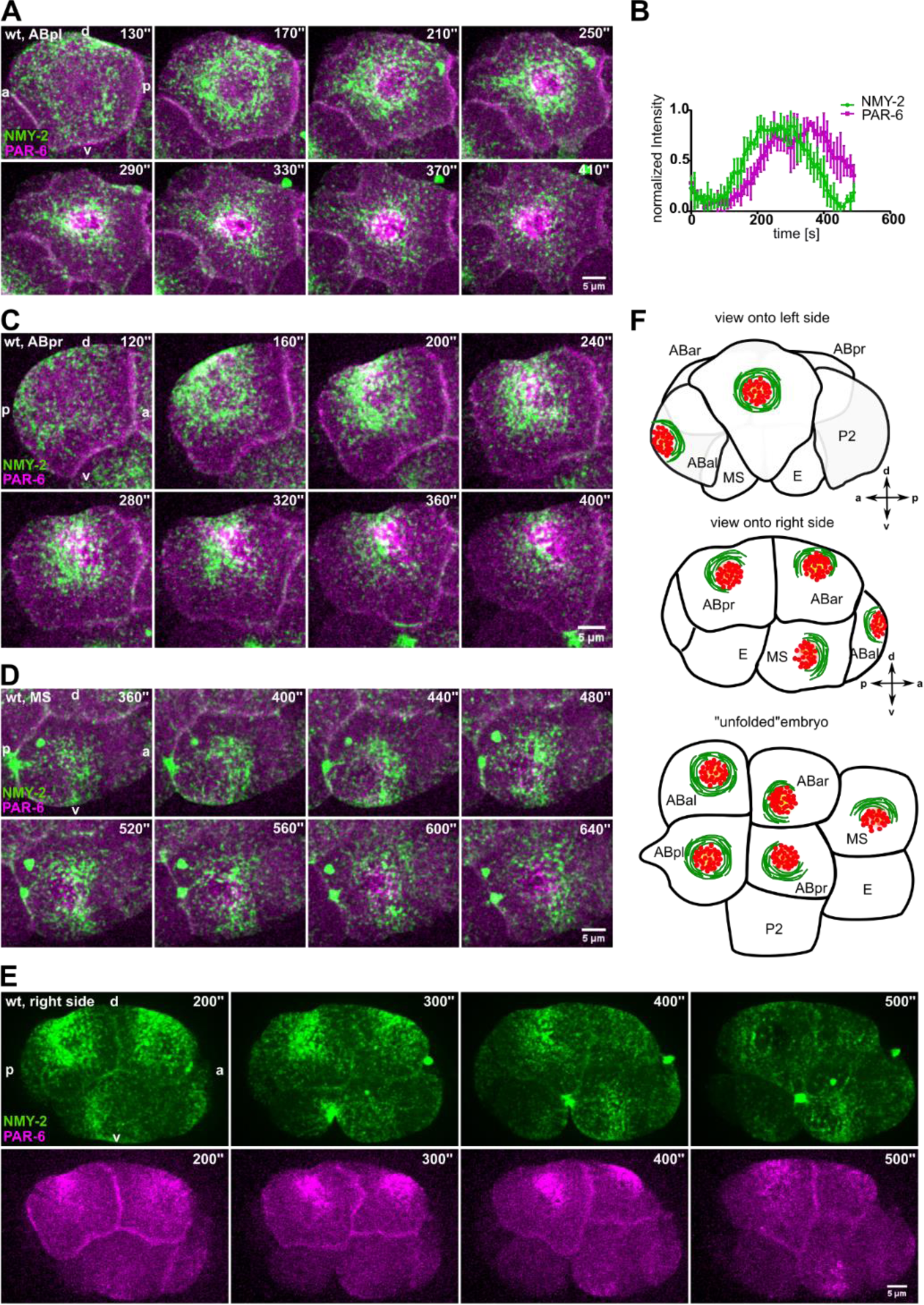
**(A, C-E)** Representative time lapse images of apical cortical sections of ABpl (A), ABpr (C), MS (D) and a right side embryo (E) expressing NMY-2::GFP and mCherry::PAR-6. Time is with respect to the completion of ABp division. **(B)** Quantification of normalized kinetics of NMY-2::GFP and mCherry::PAR-6 along time. **(F)** Top and middle: Illustration of ABpl, ABpr and MS depicting the NMY-2-aPARs domains. Bottom: Illustration depicting how an unfolded embryo would look like. The axis directions are represented below the illustration.

**Figure S2.**
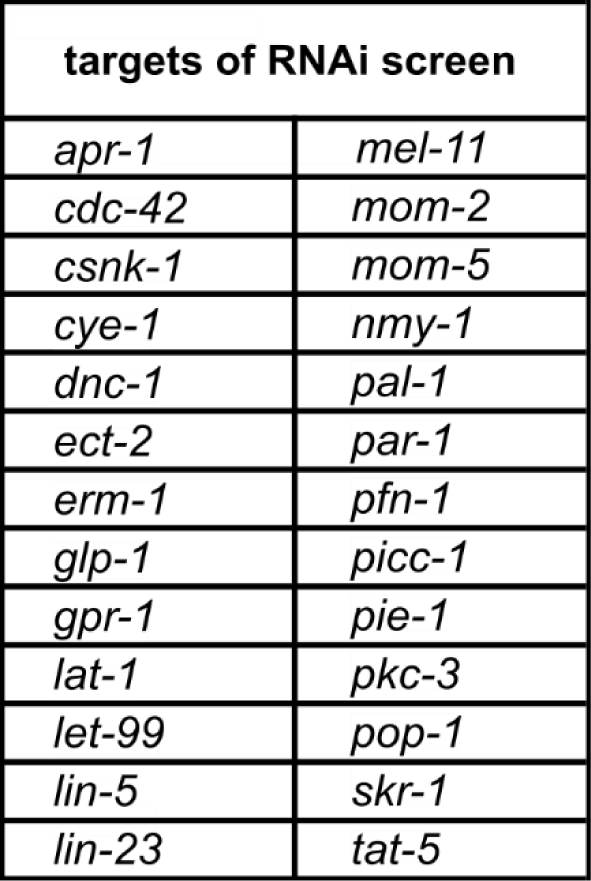
List of genes tested by targeted RNAi screening

**Figure S3.**
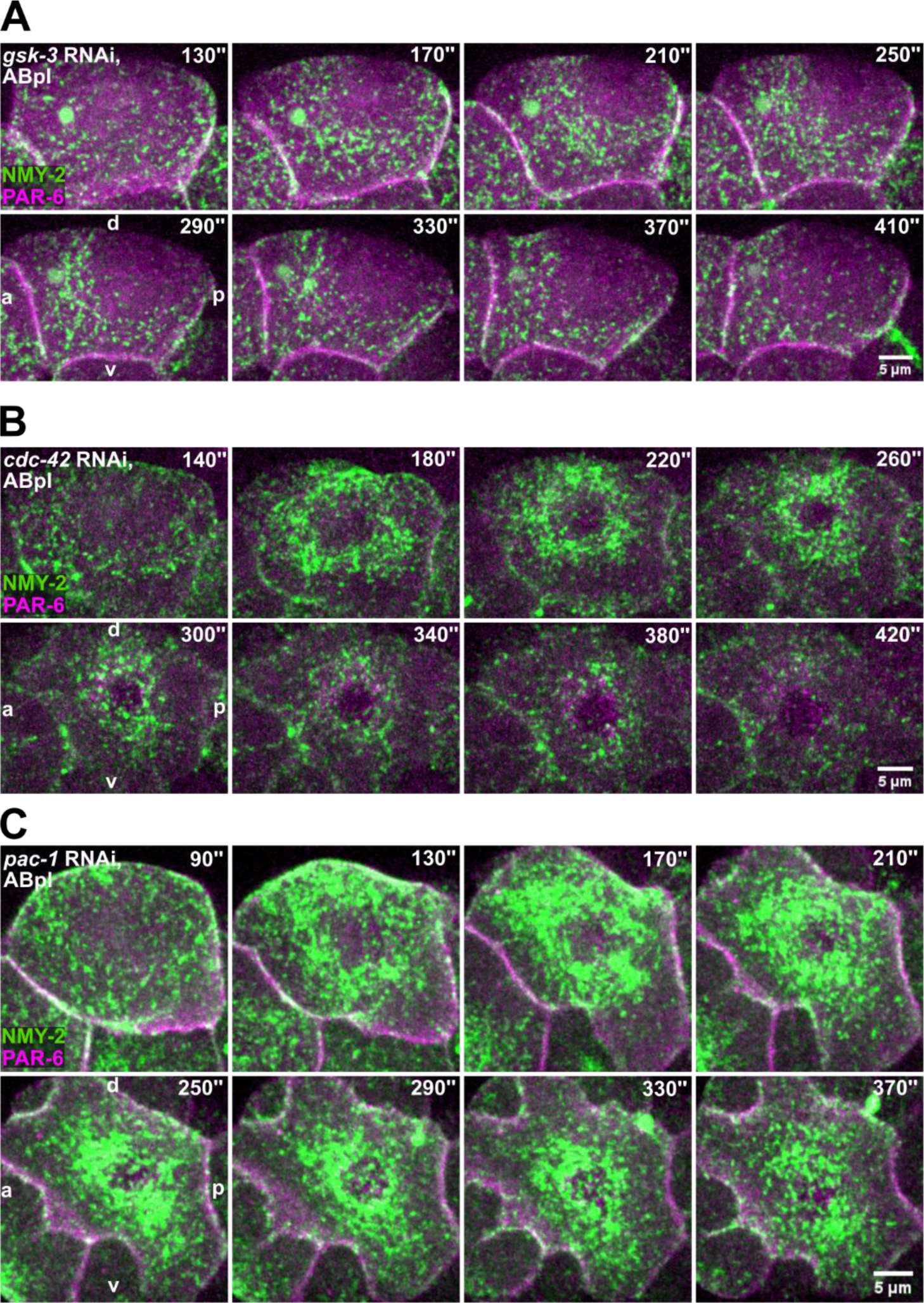
**(A-C)** Representative time lapse images of apical cortical sections of ABpl expressing NMY-2::GFP and mCherry::PAR-6 in *gsk-3* RNAi, *cdc-42* RNAi and *pac-1(xn6)* animals. Time is with respect to the completion of ABp division.

**Figure S4.**
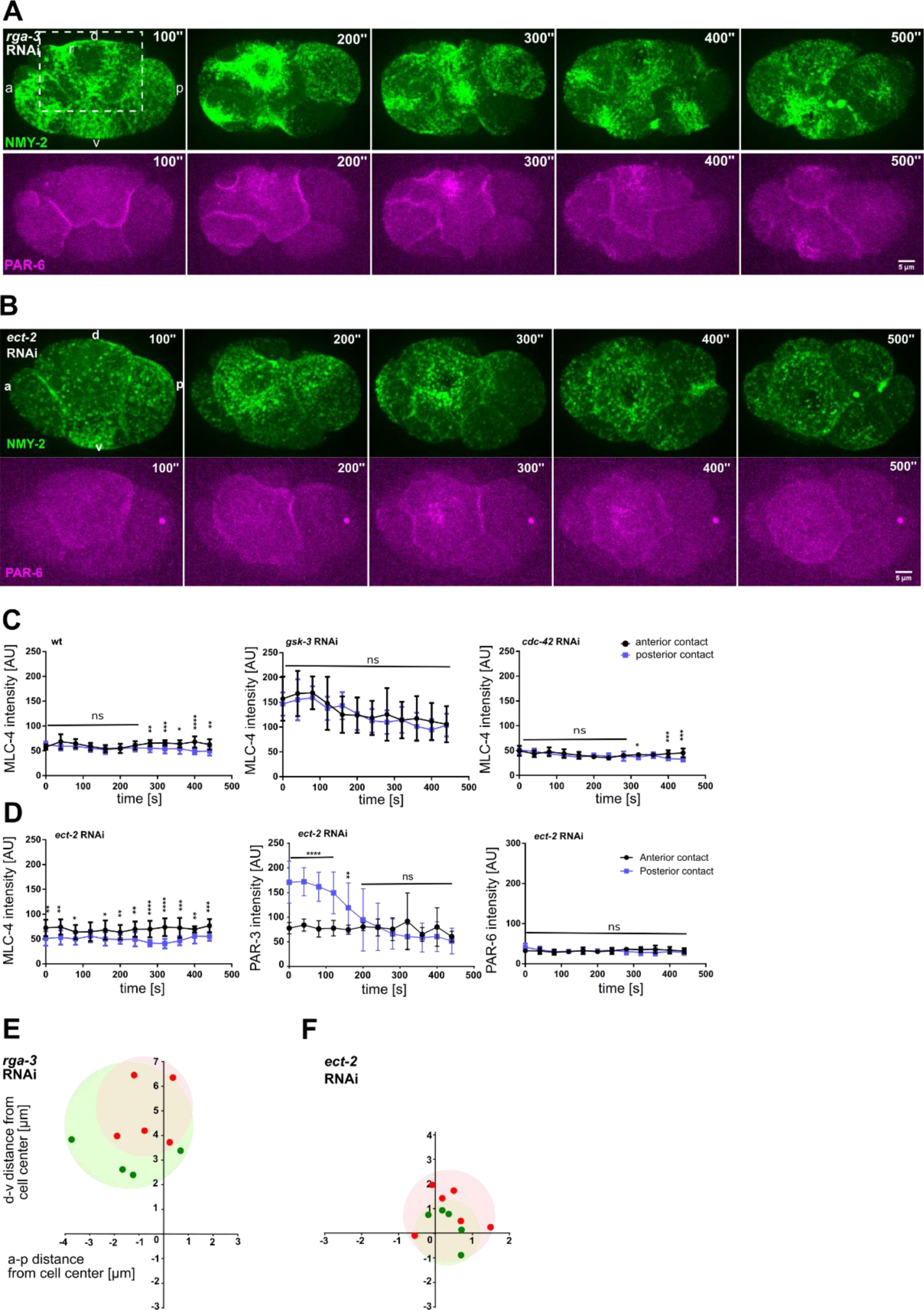
**(A, B)** Representative time lapse images of apical cortical sections of ABpl expressing NMY-2::GFP and mCherry::PAR-6 in *rga-3* RNAi and *ect-2* RNAi conditions. **(C, D)** Positioning of the NMY-2-aPAR cortical domain in *rga-3* RNAi and *ect-2* RNAi embryos with respect to the center of mass of the ABpl cell cortex which is taken as coordinate (0,0).

**Figure S5.**
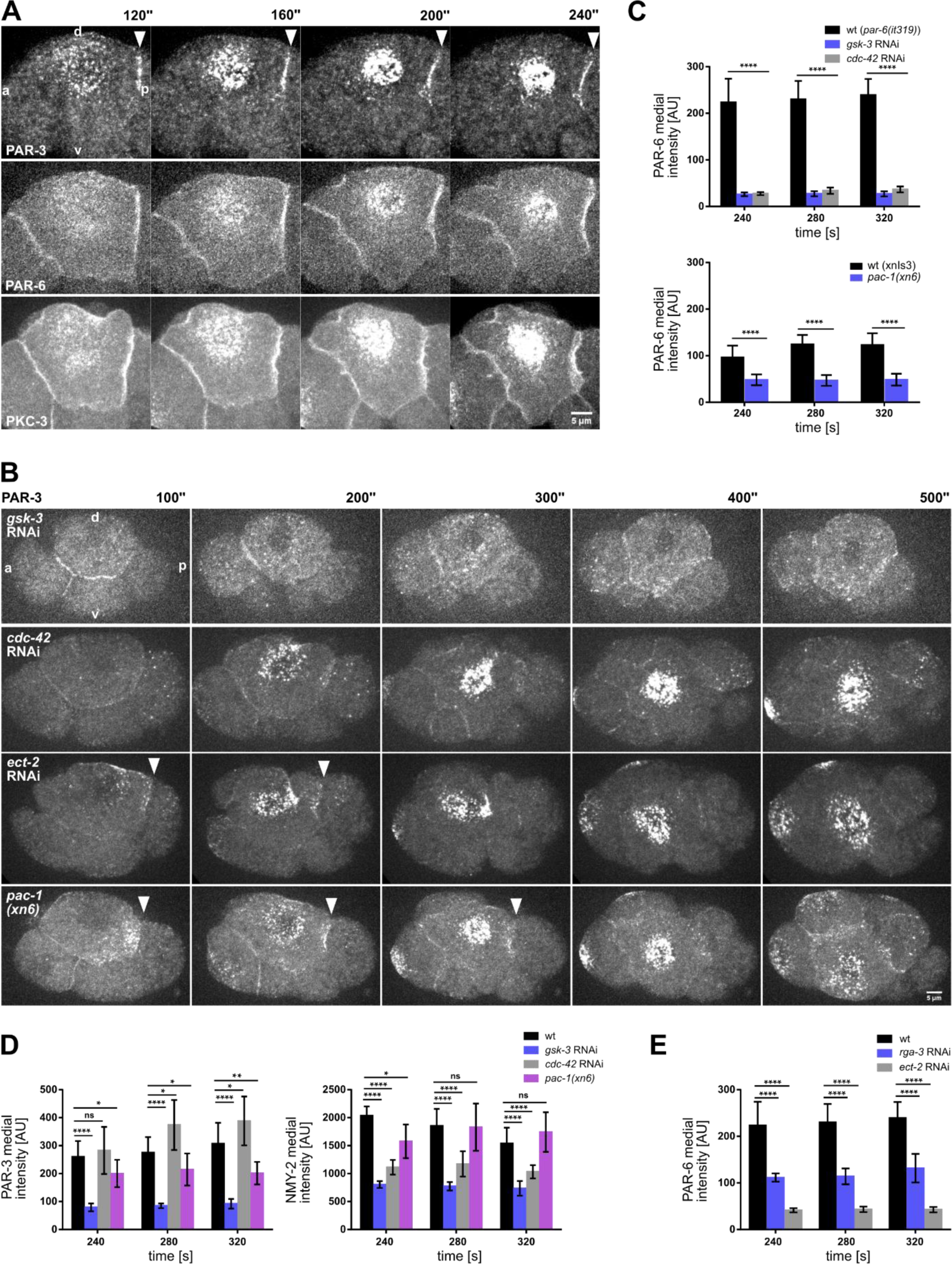
**(A)** Representative time lapse images of ABpl apical cortical sections expressing mCherry::PAR-6 (top), GFP::PKC-3 (middle) and PAR-3::GFP (bottom). Time is with respect to the completion of ABp division. White arrowheads represent asymmetric PAR-3 localization. **(B)** Representative time lapse images of cortical sections of embryos expressing PAR-3::GFP in wt, *gsk-3* RNAi, *cdc-42* RNAi, *ect-2* RNAi and *pac-1(xn6)*. White arrowheads represent asymmetric PAR-3 localization. **(C, E)** Quantification of mCherry::PAR-6 fluorescence intensity in the medial domain in RNAi conditions (n = 3). **(D)** Left: Quantification of PAR-3::GFP fluorescence intensity in the medial domain in RNAi conditions (n = 3). Right: Quantification of NMY-2::GFP fluorescence intensity in the medial domain in RNAi conditions (n = 3).

**Figure S6.**
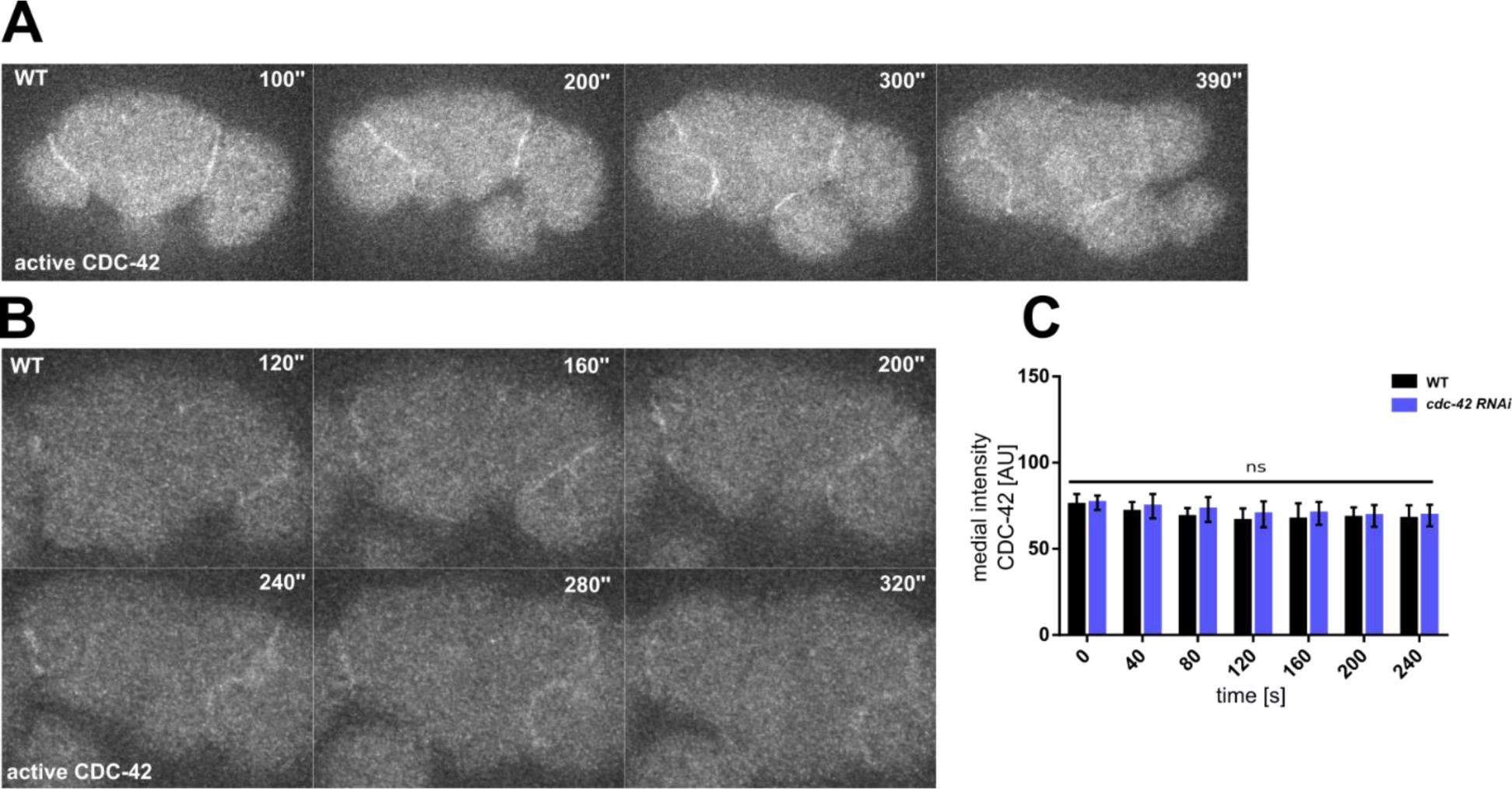
**(A-B)** Representative time lapse images of cortical sections of an embryo expressing a GFP-tagged CRIB/G-protein binding domain of WSP-1, a sensor for activated CDC-42. Time is with respect to the completion of ABp division. Panel (B) shows magnified views of ABpl. **(C)** Comparison of medial fluorescence intensity of activated CDC-42 in wt and *cdc-42* RNAi embryos.

**Figure S7.**
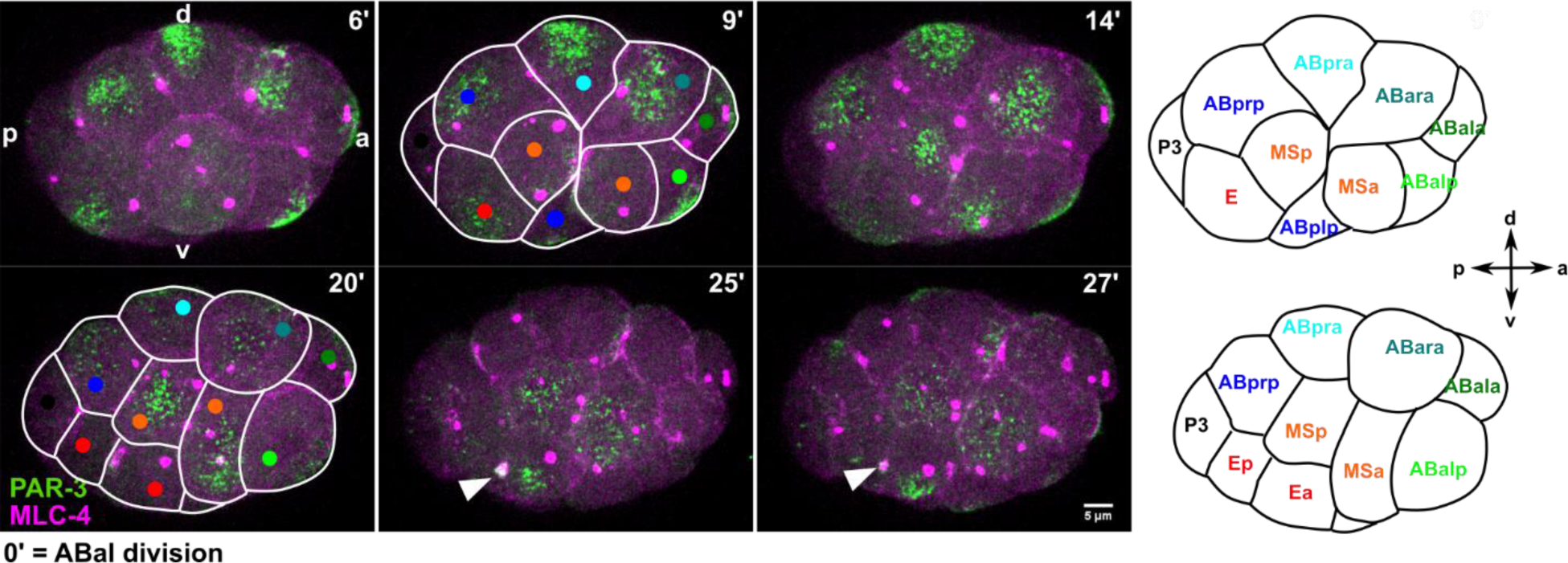
Representative time lapse images of cortical sections of embryos expressing PAR-3::GFP and mCherry::MLC-4 starting at the 12 cell stage until shortly before Ea/Ep gastrulate. White outlines depict cells intercalating into the MS furrow; white arrowhead marks the colocalization of PAR-3 with the E midbody.

### Supplementary videos

**Video S1.**

Representative time lapse series showing a wt embryo expressing NMY-2::GFP and mCherry::PAR-6. Time is with respect to the completion of ABp division. Shown is the left side.

**Video S2.**

Representative time lapse series showing a wt embryo expressing NMY-2::GFP and mCherry::PAR-6. Time is with respect to the completion of ABp division. Shown is the right side.

**Video S3.**

Representative time lapse series showing a *gsk-3* RNAi embryo expressing NMY-2::GFP and mCherry::PAR-6. Time is with respect to the completion of ABp division. Shown is the left side.

**Video S4.**

Representative time lapse series showing a *cdc-42* RNAi embryo expressing NMY-2::GFP and mCherry::PAR-6. Time is with respect to the completion of ABp division. Shown is the left side.

**Video S5.**

Representative time lapse series showing a *pac-1(xn6)* embryo expressing NMY-2::GFP and mCherry::PAR-6. Time is with respect to the completion of ABp division. Shown is the left side.

**Video S6.**

Representative time lapse series showing a *rga-3* RNAi embryo expressing NMY-2::GFP and mCherry::PAR-6. Time is with respect to the completion of ABp division. Shown is the left side.

**Video S7.**

Representative time lapse series showing an *ect-2* RNAi embryo expressing NMY-2::GFP and mCherry::PAR-6. Time is with respect to the completion of ABp division. Shown is the left side.

**Video S8.**

Combined representative time lapse series showing wt embryos expressing mCherry::PAR-6 (left), GFP::PKC-3 (middle) and PAR-3::GFP (right). Time is with respect to the completion of ABp division. Shown is the left side of embryos.

**Video S9.**

Combined representative time lapse series showing wt embryos expressing PAR-3::GFP, left side of an embryos on the left and a right side on the right. Time is with respect to the completion of ABp division.

**Video S10.**

Combined representative time lapse series showing wt, *gsk-3* RNAi, *cdc-42* RNAi and *pac-1(xn6)* embryos expressing PAR-3::GFP. Time is with respect to the completion of ABp division. Shown is the left side of embryos.

**Video S11.**

Representative time lapse series showing a wt embryo expressing PAR-3::GFP and mCherry::MLC-4 starting at the 12 cell stage until shortly before Ea/Ep gastrulate. Shown is the right side.

**Video S12.**

Representative time lapse series showing a wt embryo expressing PAR-3::GFP starting at the 12 cell stage until shortly before C divides. Shown is the left side.

### Supplementary tables

**Table S1.**

Precise timing of the three phases of cortical flow during chiral morphogenesis based on kinematic classification.

**Table S2.**

Genotypes of C. elegans strains used in this study.

